# Division and Spreading of Attention across Colour

**DOI:** 10.1101/2023.01.22.525095

**Authors:** Jasna Martinovic, Antoniya Boyanova, Søren K. Andersen

## Abstract

Biological systems must allocate limited perceptual resources to relevant elements in their environment. This often requires simultaneous selection of multiple elements from the same feature dimension (e.g., colour). To establish the determinants of divided attentional selection of colour, we conducted an experiment that used multicoloured displays with four overlapping random dot kinematograms that differed only in hue. We manipulated (1) requirement to focus attention to a single colour or divide it between two colours; (2) distances of distractor hues from target hues in a perceptual colour space. We conducted a behavioural and an electroencephalographic experiment, in which each colour was tagged by a specific flicker frequency and driving its own steady-state visual evoked potential. Behavioural and neural indices of attention showed several major consistencies. Concurrent selection halved the neural signature of target enhancement observed for single targets, consistent with an approximately equal division of limited resources between two hue-selective foci. Distractors interfered with behavioural performance in a context-dependent fashion but their effects were asymmetric, indicating that perceptual distance did not adequately capture attentional distance. These asymmetries point towards an important role of higher-level mechanisms such as categorisation and grouping-by-colour in determining the efficiency of attentional allocation in complex, multi-coloured scenes.

---

Biological systems need to allocate limited processing resources to relevant elements in their environment. Furthermore, many situations require dividing attention amongst multiple feature values: foraging for berries that could range in colour between pink, red and dark purple, or monitoring the actions of multiple players in differently coloured jerseys as well as the movement of the ball in a fast-paced team sport. In such cases, we can accept that the effectiveness of attentional selection will depend upon the relative positions of attended features within their feature space. For example, if opposing teams wore similar colours, this would make the game difficult to follow.

In vision, selection is achieved through spatial (Posner et al. 1980; Eriksen and St. James 1986), feature-based (Treue and Martinez-Trujillo 1999) or object-based attention (Duncan 1984). It is widely accepted that spatial attention operates by enhancing processing in a confined region of space through application of an attentional focus that has been likened to a spotlight, zoom lens or a Gaussian gradient (Carrasco 2014). Yet, despite decades of research, we still lack a comprehensive model that fully characterises selection of specific feature values belonging to a continuous domain of a complex, multi-dimensional feature-space such as colour (although see Stormer and Alvarez 2014).

Selection of multiple feature values from the same dimension seems highly consistent between colour (Andersen et al. 2013) and space (Toffanin et al. 2009; Adamian et al. 2019; Adamian and Andersen 2022), with reduced attentional modulation in divided attention conditions as proposed by the original Zoomlens model (Eriksen and St. James 1986) and processing of distractors modulated by target-distractor proximity in the feature-dimension on which selection is based (Treue and Martinez-Trujillo 1999; Martinez-Trujillo and Treue 2004; Martinovic et al. 2018). According to Liu and Jigo (2017), divided attention to colour is implemented by attending only a single colour at a time, which may be due to an inability to maintain multiple active attentional templates for colour. This leads to a very clear prediction that attentional modulation in the visual cortices should be halved for each individual target colour if attentional switching between the two target templates occurs in a way that ensures that both are afforded equal resources. Such a pattern has recently been reported for division of spatial attention in a multiple-object tracing task (Adamian & Andersen, 2022).

Here, we ask the following main question: how does feature-based attention operate when it has to select single or dual targets drawn from the opposite sides of the colour space? This question emerges from our recent neuroscientific work on concurrent selection of multiple hues (Martinovic *et al*. 2018), which demonstrated that different spatially overlaid colours can be attended concurrently with an efficiency that is determined by their configuration in colour space. The magnitude of attentional modulation depended overwhelmingly on target proximity and was well described by a simple model which suggested that colour space is rescaled in an adaptive manner to support attentional selection. Such adaptive rescaling is consistent with a series of recent studies that show attention optimally biases neural responses depending on several factors: the tuning of sensory neurons, physical characteristics of the visual environment and the nature of the task (Scolari and Serences 2009; Scolari et al. 2012; Verghese et al. 2012). It is also consistent with the recent proposals integrating various evidence streams from the visual search literature (Geng and Witkowski 2019; Yu et al. 2023): attentional guidance only has to be ‘good-enough’ to efficiently direct the observer towards the best approximation of the target in the current sensory context, which is why it is highly dependent on the configuration of targets and distractors in feature space.

Whilst Martinovic et al. (2018) examined the behavioural and neural signature of attentional selection for different sets of dual targets in two different colour configurations, the key condition of attending single colour targets in the same contexts was missing. Evaluating differences between single target and two-target attentional settings is essential in an attempt to understand the effectiveness of attending to multiple targets (e.g. Liu and Jigo 2017, for a review see Ort and Olivers 2020). Therefore, to address this gap in the literature, we examined the behavioural and neural outcomes of applying single or dual attentional foci in a task that requires sustained feature-based selection. To fully assess how feature-based attentional selection works for individual colours in a display with multiple targets and distractors, it is optimal to be able to record an individual response for each colour. We achieved this by using a multi-colour random dot kinematogram (RDK) display in combination with frequency-tagging, allowing us to look at the neural markers of attentional processing for 4 colours simultaneously. This is a well-established procedure for studying the neural markers of feature-based attention (e.g., Andersen and Muller 2010; Andersen, Fuchs, et al. 2011; Andersen et al. 2012; Andersen *et al*. 2013; Andersen et al. 2015; Gundlach, Forschack, et al. 2023; Gundlach, Wehle, et al. 2023). The colours differed in hue but were of constant lightness and colourfulness, as specified in a perceptually uniform colour space (CIE Lab; see Fairchild 2013).

We measured neural activity elicited by each colour in a 4-colour display when attention was focused on red or green or divided between red and green in order to detect infrequent coherent motion events in these colours. To generate different stimulus contexts in which these targets would need to be selected, we manipulated distances between targets and one of the distractor hues. This resulted in three contexts: i) a putatively neutral context, with intermediate distractors falling between the two target hues (blue and yellow); ii) a context with one distractor closer to red (orange and yellow); and iii) a context with one distractor closer to green (lime and yellow). We measured neural and behavioural outcomes of attention to single or multiple colours using the same paradigm with and without flicker. If application of multiple attentional foci in colour space divides the processing resources equally between the two targets, then attentional modulation of SSVEPs elicited by targets should be halved when moving from single to dual targets. Increases in target-distractor proximity should also incur costs in terms of decreased effectiveness of selection – behaviourally, by reduced sensitivity to signal colour; neurophysiologically, by SSVEP amplitudes driven by targets being decreased and SSVEP amplitudes driven by distractors being increased. Finally, if operational characteristics of perception and attention are both well captured within the same perceptually uniform colour space, equal distance in colour space should yield equal effectiveness of attentional selection.

## Materials and Methods

### Participants

Twenty-five volunteers in total took part in the study, but five were removed from the final sample. One participant was excluded due to technical problems during the EEG recording. Four other participants were excluded due to inadequate task performance in the EEG session (identified as d’s < 0 in any of the conditions). The remaining 20 participants (4 male) were on average 23 years old (range 21–27 years). With a sample size of 20 participants and a significance level of 0.05, a paired samples t-test would yield 80% power with an effect size of d=0.66 (calculated with package pwr for R; Champely et al. 2020). In light of previous work in the field, which generally reports large effects of attention on SSVEP amplitudes (>0.8; e.g. Andersen et al. 2008; Andersen *et al*. 2012) this was deemed to be a sufficiently large sample.

All participants were right-handed and had normal or corrected-to-normal vision and normal colour vision as assessed by the Cambridge Colour vision test (CCT; Regan et al. 1994). All participants reported having no neurological or psychiatric history and gave written informed consent prior to testing. Participants were recruited amongst the University of Aberdeen students and were reimbursed for their time and effort. The study was approved by the ethics committee of the School of Psychology, University of Aberdeen and was in line with the Declaration of Helsinki (1964).

### Stimuli and Procedure

Stimuli were presented on a 21-inch ViewSonic P227f CRT monitor, calibrated using a ColorCAL 2 (CRS, UK) and controlled by a VISaGe (CRS, UK) system. The resolution was set to 640 x 480 pixels with a refresh rate of 120 Hz. CRS toolbox and CRS colour toolbox for Matlab (Mathworks, USA) were used to run the experiment. Colour conversions were based on 1931 colour matching functions and measurements of the spectra of the monitor phosphors taken by a SpectroCAL (CRS, UK). The behavioural session was done in a dark room and the EEG session in an electrically shielded, sound-attenuated chamber, each with a ViewSonic P227f monitor controlled by a VISaGe system. The viewing distance was approximately 70 cm. The monitor was the only source of light.

The stimuli consisted of four fully overlapping random dot kinematograms (RDKs) that differed in hue (0°, 180°, 90°, 270°, 45° and 135°, corresponding to red, green, yellow, blue, orange and lime; see Table 1) but were of equal chroma and lightness in CIE LCh space (C=34 and L=60). Perceptual colour spaces are continuous 3D spaces with the dimensions of hue, colourfulness and lightness (e.g., CIELAB, CIELUV; Fairchild 2013). Euclidean distance (ΔE) between two points in these spaces is taken to be a measure of their perceptual colour difference, with a just-noticeable difference being equal to a ΔE of 2.3. CIELAB space is used in studies on colour-based attention under the assumption that it is fully perceptually uniform and therefore enables researchers to collapse data across individual colours, as long as same colour distances between targets and distractors are used (e.g. Stormer and Alvarez 2014). However, while CIELAB does a decent job at representing appearance of colours, it is worth noting that the perceptual unique hues (red, green, yellow and blue) do not align directly with the CIELAB axes (Fairchild 2013). Assuming illumination equivalent to D65, unique hues lie approximately at CIELAB hue angles of 24° (red), 90° (yellow), 162° (green), and 246° (blue) (Fairchild 1996; for an in-depth analysis, see Seymour 2020; Figure 1a). Despite these caveats, CIELAB is well suited for the current study because it provides a space in which relative perceptual distances between hues can be easily controlled, while ensuring that saturation, colourfulness and lightness remain constant across samples (Schiller et al. 2018).

**Figure 1.**
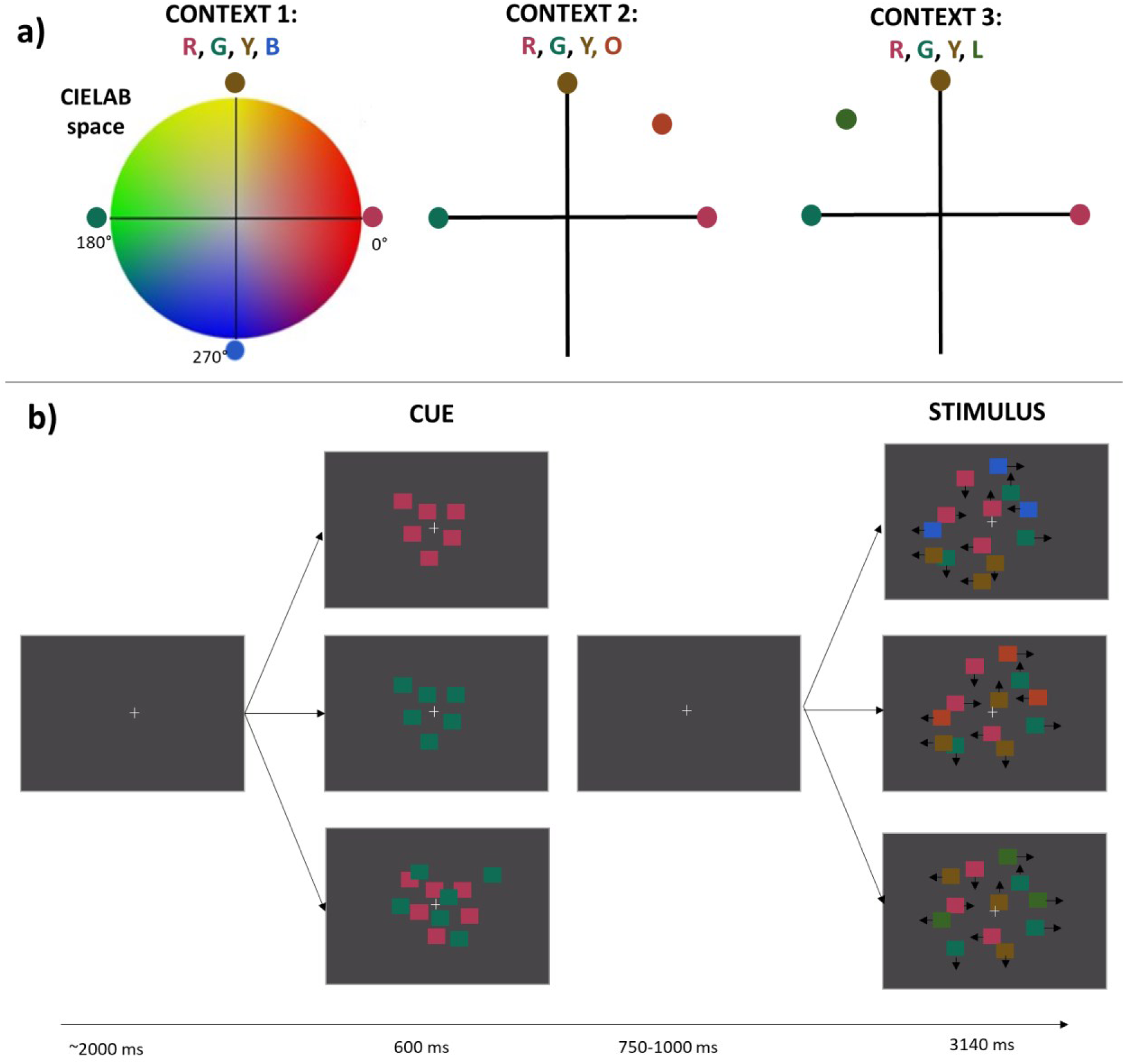
Stimulus colours and trial outlook. A) Stimulus colours. Colours were selected from the perceptually uniform CIELAB space (see *Table 1* for coordinates) to generate three sets of contexts. Red and Green were targets in each context. Yellow was the fixed distractor, also present in each context. There were three context-specific distractor colours: 1) blue, generating a context with equally spaced colours in CIELAB; 2) orange, generating a context in which a distractor was closer to the red target; and 3) lime, generating a context in which a distractor was closer to the green target. B) Trial outlook. Participants were cued to attend either red or green alone or both colours simultaneously. After a delay of 0.75–1.0 s they observed a stimulus interval with the four colour RDKs intermingled spatially and moving at random. Each stimulus interval could contain between 0 and 3 brief coherent movements (left, right, up or down) of either the attended or the unattended colours. In the figure, the number of dots is reduced for the sake of clarity. Colours cannot appear exactly as they did on a calibrated display but have been adjusted to closely resemble their appearance in sRGB space.

**Table 1.** Stimulus and background colour coordinates in CIELCh and CIE 1931.

| Colour | CIELCh |  |  | CIE 1931 |  |  |
| --- | --- | --- | --- | --- | --- | --- |
|  | L | C | h | x | y | Y |
| Red | 60 | 34 | 0 | 0.380 | 0.297 | 28.12 |
| Green | 60 | 34 | 180 | 0.247 | 0.361 | 28.12 |
| Yellow | 60 | 34 | 90 | 0.397 | 0.418 | 28.12 |
| Blue | 60 | 34 | 270 | 0.230 | 0.242 | 28.12 |
| Orange | 60 | 34 | 45 | 0.425 | 0.361 | 28.12 |
| Lime | 60 | 34 | 135 | 0.322 | 0.426 | 28.12 |
| Background | 55 | 0 | 0 | 0.313 | 0.329 | 23.00 |

The approximate appearance of the stimuli is depicted in Figure 1, together with the trial outlook. Colours were presented in three colour contexts. In the first context, the colours were equally spread in CIELAB space (red, yellow, green and blue). In the second context, we kept the yellow distractor and replaced the blue distractor with orange, to place it closer to red (red, orange, yellow and green). Lastly, in the third context, the blue distractor was replaced with lime, to place it closer to green (red, yellow, lime and green). Rather than deploying distractors and targets pooled from different sections of the CIELAB as in Stormer and Alvarez (2014), we opted to evaluate in detail shifts within one quadrant of the colour space. This allows us to quantify any hue-specific non-uniformities. Within our design, the yellow distractor served as an important comparison point both between and within colour contexts. Meanwhile, the orange and lime distractors are both placed in the same half of hue space but occupy a position nearer to one of the targets (red and green, respectively). The background was metameric to D65 (CIE 1931 coordinates: x=0.3128, y=0.3290, Y=23 cd/m^2^).

For the EEG session each colour was tagged with a specific flicker frequency. The colour red flickered at 8.57 Hz; green at 10 Hz; yellow at 12 Hz and depending on the context either blue, orange or lime at 15 Hz. These frequencies correspond to flicker cycles of 14, 12, 10, and 8 frames, respectively, at our monitor refresh rate of 120 Hz. The on-off ratio for each frequency was 50-50; e.g., 10 Hz flicker was produced by presenting the dots for 6 frames and then switching them off for another 6 frames.

While luminance and chromatic mechanisms have different temporal response dynamics, SSVEPs driven by combined colour and luminance signals are an outcome of non-linear summation of the two constituent signals which combines properties of both, with a faster latency than an isoluminant chromatic stimulus but also a higher amplitude than a luminance-isolating stimulus (Martinovic and Andersen 2018). SSVEPs driven by different cone-opponent signals sum differently with luminance contrast, and this summation is also likely to be influenced by temporal frequency. However, when selective attention is deployed to one of two colours, attentional modulation of the said SSVEP signal remains rather stable across different hues, as long as they are all superimposed on equivalent luminance pedestals (Martinovic and Andersen tested 0.37 and 0.71 Weber contrast; in the present study, the luminance pedestal equates to 0.223 Weber contrast). This makes the SSVEP frequency tagging method highly suitable for studying colour-based attention. However, the presence of luminance contrast in the stimulus should not be fully neglected when interpreting the results, as it facilitates the performance of the coherent motion detection task. In isoluminant conditions, the coherent motion task would become more difficult and in the presence of relatively fast flicker would likely become even more driven by the L and M-cone related signals picked up by the luminance system (for potential models, see Mullen et al. 2003; Allard and Faubert 2014). In the present study, attentional selection itself remains orthogonal to motion coherence processing: attention needs to target the chromatic component of the stimulus since the luminance component is constant and thus uninformative for target identity (for an extended discussion, see Martinovic & Andersen, 2018).

Final methodological consideration concerns the use of a fixed assignment of flicker frequencies to colours, rather than a counterbalanced assignment. Although the absolute magnitude of SSVEP amplitudes differs across colours and across frequencies, we have generally observed equivalent patterns of attentional modulation of colour for frequencies in the range employed here (Adamian and Andersen 2023). Thus, any potential benefit of counterbalancing frequencies would not have outweighed the costs in terms of complexity of analysis and reduced precision of measurement, due to the increased number of conditions for each colour, but with fewer trials per condition.

The field occupied by the random dot stimuli extended over a circular area with a diameter of 15.642°. Each of the four colours was represented by 75 squares of 0.391° of visual angle, which made 0.061° displacements per frame in a random direction. The squares were drawn in random order to prevent systematic overlap. At the start of each trial a white fixation cross (0.488° by 0.488°) was presented for about 2 seconds, followed by the presentation of a cue that lasted 600 ms. The cue instructed participants to attend to one colour (either red or green) or two colours (both red and green), by presenting a static frame of dots in the to-be-attended colour(s). The cue was followed by 0.75-1 seconds of fixation prior to the motion interval with all four RDKs, which lasted 3.14 seconds. The motion display also contained the fixation cross and participants were instructed to maintain their gaze at fixation throughout each trial. Target and distractor coherent motion events were embedded within Brownian-like dot motion, with the first 500 ms being event-free. Participants used a CT-6 button box (CRS, UK) and were instructed to respond to coherent motion of the cued colour(s), whilst ignoring any such motion from uncued colours. A coherent motion lasted 400 ms and consisted of 50% of the dots from the same colour, moving in the same cardinal direction (up, down, left or right). Dots that constituted the 50% coherence motion event were randomly selected on each frame to further prevent any spatial tracking strategy from being effective. The onsets of sequential coherent motion events were separated by at least 700 ms. Responses that occurred within 250-900 ms after the onset of coherent motion events in attended or unattended dots were counted as hits or false alarms, respectively. Similarly, events that did not receive a response in this period were counted as misses or correct rejections. As the task is to attend a specific colour for the purpose of coherent motion detection (i.e., participants must select the correct target colour to be able to perform the task accurately, once they detect a motion signal), making a false alarm to a distractor-colour coherent motion or missing a target-colour coherent motion is driven by a lack of sensitivity for colour, and thus aligned with the assumptions of the signal detection theory. Participants undergo training prior to the experiment to learn to perform the coherent motion task at ceiling level. Any responses to non-coherent motion would result from noise crossing the decision criterion in the motion signal detection system. Such responses are extremely rare, as participants quickly learn to respond only to coherent motion.

The experiment consisted of two separate sessions: a behavioural and an EEG session. The main differences between them were (1) the inclusion of flicker-frequency tagging and (2) the reduction in the number of coherent motion events in the EEG session, aimed at effectively capturing attention-related processing using SSVEPs (for a review, see Andersen, Müller, et al. 2011). Before the start of the behavioural session, participants were asked to do a practice block of 40 trials. During practice they were provided with feedback sounds: a low beep (600 Hz) corresponded to an incorrect response (false alarm/miss) and a high beep (1500 Hz) indicated a correct response (hit). Prior to the EEG recording, participants did another practice set of 15 trials to familiarise themselves with the now flickering stimuli.

For the behavioural session, participants performed 270 trials, with 30 trials for each of the nine conditions (3 attentional conditions x 3 colour contexts). All the trials contained between 1 and 3 coherent motion events, resulting in 540 events in total and 60 events per condition (15 per colour). This means that each participant was presented with 180 target events (90 per colour), 90 red and green distractors (45 per colour), 135 yellow distractors, and 135 blue/lime/orange distractors (45 per colour). In the EEG session there were a total of 405 trials (45 per condition). In each condition, 30 trials contained no events and 15 trials had between 1 and 3 events, for a total of 30 events (i.e., 270 events across the whole experiment). It was not possible to equalise the number of target and distractor events across colours within these parameters so these were assigned randomly, resulting in small differences between the number of events in each colour seen by each of the participant (ranges for red and green targets: 81 – 103 events, red and green as distractors: 35-53, yellow distractors: 53-76, lime/blue/orange distractors: 58 – 73). These small fluctuations are fully accounted for within the statistical analysis, as generalised linear mixed effect statistical models are fit to the data combined across experiments with experiment as a fixed effect and participant as a random effect.

The completion of the behavioural session, including the colour vision test and one practice block took approximately an hour. The EEG session lasted about 2½ hours, with the recording lasting approximately 1 hour and the rest of the time devoted to the brief practice block and electrode setup and removal.

### Experimental Design and Statistical Analysis

#### Behavioural data

Statistical analyses were implemented in R (version 4.2,1, R_Core_Team 2016), using packages tidyverse (version 1.2.1, Wickham et al. 2019), lme4 (version 1.1-31, Bates et al., 2015), effectsize (version 0.8.2, Ben-Shachar et al., 2020), emmeans (version 1.8.3, Lenth et al., 2019), performance (version 0.10.1, Lüdecke et al. 2021), DHARMa (version 0.4.6, Hartig 2022) and lmeresampler (version 0.4.2, Loy 2023).

To establish how attentional focus and stimulus context affected participant performance, we analysed dichotomous outcomes elicited by targets and distractors (i.e. whether events were responded or not responded to). We used generalised linear mixed effect models (GLMMs) on the binomial single-trial data elicited by target or distractor colours, as implemented in the R statistic package lme4. While ordinary logit models have many advantages over ANOVAs on percentage data (e.g. they resolve the potential issue of confidence intervals outside of the 0-100% range), mixed logit models have the further advantage of being able to account for random subject effects (Jaeger 2008). As mentioned earlier, we combined the behavioural data from both experiments prior to fitting the models. Contrasts were set to simple coding: each level of the factor was compared to the reference level, with the intercept set to grand mean. Full details of the best fitting model, including its reference levels, is presented in Supplementary Materials. When fitting GLMMs, maximal random effect structure that was possible was applied, while maintaining goodness of fit (Barr et al., 2013). We verified that the model residuals were normally distributed and did not suffer from overdispersion or excessive outliers using a simulation-based approach implemented in the DHARMa package (Hartig 2022). We also verified that models with multiple predictors did not suffer from predictor collinearity using the check_colinearity function from the *performance* R package on an additive model (Lüdecke *et al*. 2021).

Models were statistically evaluated by iteratively removing higher-order interactions between fixed effects from the model to verify if it significantly reduced their goodness of fit. In Supplementary Materials we report all estimates of the best-fitting, final model, including all the fixed effects and interactions that could not be removed. Post-hoc tests on this final model were performed using omnibus paired t-tests on the highest order interactions remaining in the model, corrected for multiple comparisons (p<.05) with the ‘mvt’ method from emmeans package (Lenth et al., 2019). This method relies on the multivariate t distribution with the same covariance structure as the estimates to determine the p-value adjustment.

Responses to targets were fitted with a GLMM with fixed effects of *context (blue, orange or lime), attentional focus (focused on a single target or divided between two targets)*, *target colour eliciting the response (red or green)* and *experiment (behavioural or EEG).* Responses to distractors were analysed separately for the fixed distractor (yellow) and the context-dependent distractors (blue, orange and lime). For yellow, the GLMM included the fixed effects of *context (blue, orange or lime), attentional focus (to red, green or both targets)* and *experiment (behavioural or EEG).* For blue, orange and lime, the GLMM included the fixed effect of *distractor colour (blue, orange or lime)*, *attentional focus (to red, green or both targets)* and *experiment (behavioural or EEG).* Finally, when red and green were attended on their own, these two colours would also act as distractors. Responses driven by these red and green distractors were also investigated by fitting a GLMM with the fixed effects of *context (blue, orange or lime)*, *distractor colour (red or green)* and *experiment (behavioural or EEG).* This control analysis aimed to verify if the target colours were equally effective when they adopted the role of a distractor. Having selected two colours from the opposite sides of the colour space, we expected that this would be the case.

To narrow down the interpretation of target and distractor-related performance, we also analysed response biases. We used criterion (c) as the most basic measure of response bias, combining false alarms (FAs) across the two distractor colours into a single FA rate prior to the calculation of criterion for red, green or dual targets for each colour context. When false alarm and hit rates sum to 100% c = 0, if both are 99% then c = −2.33 and if both equal 1% then c = 2.33 (Macmillan and Creelman 2004). Thus, criterion below 0 reflects a liberal response pattern (FAs exceed misses) and criterion above 0 reflects a conservative response pattern (misses exceed FAs). Prior to calculating the criterion, hit rates of 100% and FA rates of 0% were adjusted according to Wickens’ (2002) proposal, which assumes that should the number of trials have been doubled, at least one hit or FA would have been observed. The criterion is informative on whether attention to multiple targets leads to a change in the position of the ‘decision axis’ about the presence or absence of the signal. The number of target events when attention is divided across two colours increased from 25% to 50% of all coherent motion events, while the number of distractor events fell from 75% to 50%. Target prevalence influences response criterion rather than sensitivity (i.e. d’) as it leads to trade-offs between misses and FAs (Wolfe and Van Wert 2010). In figure 2a depicting the results from Wolfe & Van Wert’s (2010) visual search study, the shift in criterion remains negligible when manipulating target prevalence between 25% and 50%. Nevertheless, quantifying the criterion in addition to the hit and FA rates will allow for a more balanced interpretation of the behavioural findings. We analysed criterion shifts by fitting a linear mixed effect model (LMM) with the fixed effects of *context (blue, orange or lime), attentional focus (to red, green or both targets)* and *experiment (behavioural or EEG)*.

#### EEG recording, processing and analysis

Brain electrical activity was recorded at a sampling rate of 256 Hz from 64 Ag/AgCl electrodes using a Biosemi ActiveTwo EEG system (BioSemi, Netherlands). Lateral eye movements were monitored with a bipolar outer canthus montage (horizontal electrooculogram). Vertical eye movements and blinks were monitored with a bipolar montage positioned below and above the left eye (vertical electrooculogram). During recordings, the CMS (Common Mode Sense) and DRL (Driven Right Leg) electrodes were used to drive the average potential for each participant as close as possible to the AD-box reference.

EEG data were processed using the EEGLab toolbox (Delorme & Makeig, 2004) in combination with custom-made procedures in Matlab (The Mathworks, Natick, MA). A period of 500 ms after stimulus onset was discarded to exclude the evoked response to stimulation onset and to allow the SSVEP sufficient time to build up. Further, from 2900 ms after stimulus onset, the data were also discarded, as in this period participants might become aware that no further events were possible, and this knowledge could have led participants’ attention to diminish. This resulted in epochs of 2400 ms duration being extracted for SSVEP analysis. All epochs with target or distractor onsets occurring within the epoch were excluded from the SSVEP analysis. This ensured that the analysed data were not contaminated by activity related to coherent motion or manual responses and left a total of 30 epochs for each condition. All epochs were detrended (removal of mean and linear trends). Epochs with eye movements or blinks were rejected from further analysis, and all remaining artifacts were corrected or rejected by means of an automated procedure (FASTER; Nolan et al., 2010). The average rejection rate was 12%, leaving an average of 27 trials per condition for analysis (range 14-30). While FASTER was conducted using Fz as reference, data were subsequently transformed to average reference. All epochs within the same condition were averaged for each participant. These means were Fourier-transformed, and SSVEP amplitudes were quantified as the absolute value of the complex Fourier-coefficients at the four stimulation frequencies. Based on the topographical distribution of mean SSVEP amplitudes a cluster of 5 electrodes (POz, Oz, O1, O2, Iz) where amplitudes were maximal was chosen and amplitudes were averaged across these electrodes for statistical analysis. Attentional effects were computed by subtracting amplitude when unattended from amplitude when attended (A-U).

To investigate how the attentional increase in SSVEP amplitudes was influenced by the different conditions of the experiment, we fitted linear mixed effect models (LMMs) on the strength of the attentional effect (A-U) with *attentional focus (to a single target or divided between two targets), context (blue, orange or lime)* and *target colour from which responses are analysed (red or green)* as fixed effects and participant as a random effect. We opted for LMMs as they are robust to non-normally distributed data and instead require the residuals to be normally distributed (Schielzeth et al. 2020). Since the changeable distractors were physically different (blue, orange and lime) and thus elicited vastly different amplitudes (consistent with e.g. Martinovic and Andersen 2018; Martinovic *et al*. 2018), the distractor-driven amplitudes could not be fully compared between contexts. To investigate how distractors were processed between contexts, we therefore focused exclusively on the SSVEP amplitude elicited by yellow, which was the distractor that was constantly present. A LMM analysis with *context (blue, orange or lime),* and *attentional focus (to red, green or split)* as fixed effects and participant as a random effect was conducted. We also fitted a LMM with *attentional focus (red, green or split)* as a fixed effect and participant as a random effect to amplitudes driven by each of the context-specific distractor colours (blue, orange and lime) separately, to reveal whether their neural processing was affected by the target colour(s). The LMMs were run using the same software tools as for the behavioural data, including the same method for conducting assumption checks and pairwise differences for the estimated means.

Finally, to understand the properties of the putative hue-selective foci more fully, we further evaluated differences in amplitudes between single-target and dual-target selection. Our simple model is based on selection of single or multiple confined regions in hue-space and posits that colour selection operates through modulating the processing of neurons that respond to the hue(s) within this region. Both the dimension-weighting and relational models of attention agree that attentional settings for colour are influenced by their categorical status (Found and Muller 1996), which can be used to set selection weights relative to the other colours in the display (e.g. ‘redder than’ or ‘yellower than’; Becker 2010). To capture this breadth of target-representation tuning, we assume a relatively wide selection region, encompassing an area of +/- 45° in colour space (for similar extent of colour tuning in visual working memory, see Bays et al. 2009). A simple mechanistic model, in which a transition from one to two attentional foci would reduce the processing resources in half, would predict that attentional SSVEP modulation should be halved when combining the two attentional settings. This is equivalent to evaluating a mean-amplitude model in which amplitude when selecting two features concurrently is equal to the average of amplitudes when attending single feature values individually (for similar findings from orientation and colour, see Andersen *et al*. 2008; Andersen *et al*. 2015). The mathematical equivalence of these statistical tests stems from the fact that attentional increases when attending red alone or together with green are calculated by subtracting the same unattended amplitude (i.e., red when green is attended). Thus, despite the values being different, the statistical comparison relies on the identical underlying data (i.e. testing if (AmpR_,r_ + AmpR_g_) / 2 = AmpDiv_r_ vs. (AmpR_r_ – AmpR_g_) / 2 = AmpDiv_r_ – AmpR_g_ where AmpR_r_ is amplitude for red when attending red, AmpR_g_ is amplitude for red when attending green and AmpDiv_r_ is amplitude for red when dividing attention). These predictions were evaluated by solving for the multiplier *p* in the following equations: AmpDiv_r_ = p * (AmpR_,r_ + AmpR_g_) and AmpDiv_g_ = p * (AmpG_g_ + AmpG_r_). We then compared the difference between the obtained values and 0.5, to evaluate the accuracy of our predictions.

## Results

### Behavioural findings

For the behavioural experiment, collapsing the data across all stimuli and conditions yields hit rates of 78% ± 12%, false alarm rates of 6% ± 5%, with a criterion of 0.36 ± 0.27 (means and standard deviations). In the EEG experiment, hit rates were 65% ± 13%; false alarm rates 9% ± 10% and criterion 0.43 ± 0.31 (means and standard deviations; see Figure 3). Median number of button presses occurring when no coherent motion event was presented was very low, at 3 per participant across the duration of the EEG experiment (range 0-22). This shows that the motion detection task is challenging but achievable, reflected by the high percentage of hits and the low rate of false alarms. A positive but relatively small criterion value indicates that participants adopted a somewhat conservative strategy. Once again this confirms that the task was achievable. The small decrease in overall hit rates and increase in false alarm rates in the EEG experiment can be attributed to the addition of flicker. Some participants particularly struggled performing the motion detection task on flickering RDKs: as described in the Participant section, 4 observers became unable to perform the task reliably and were removed from the sample.

**Figure 3.**
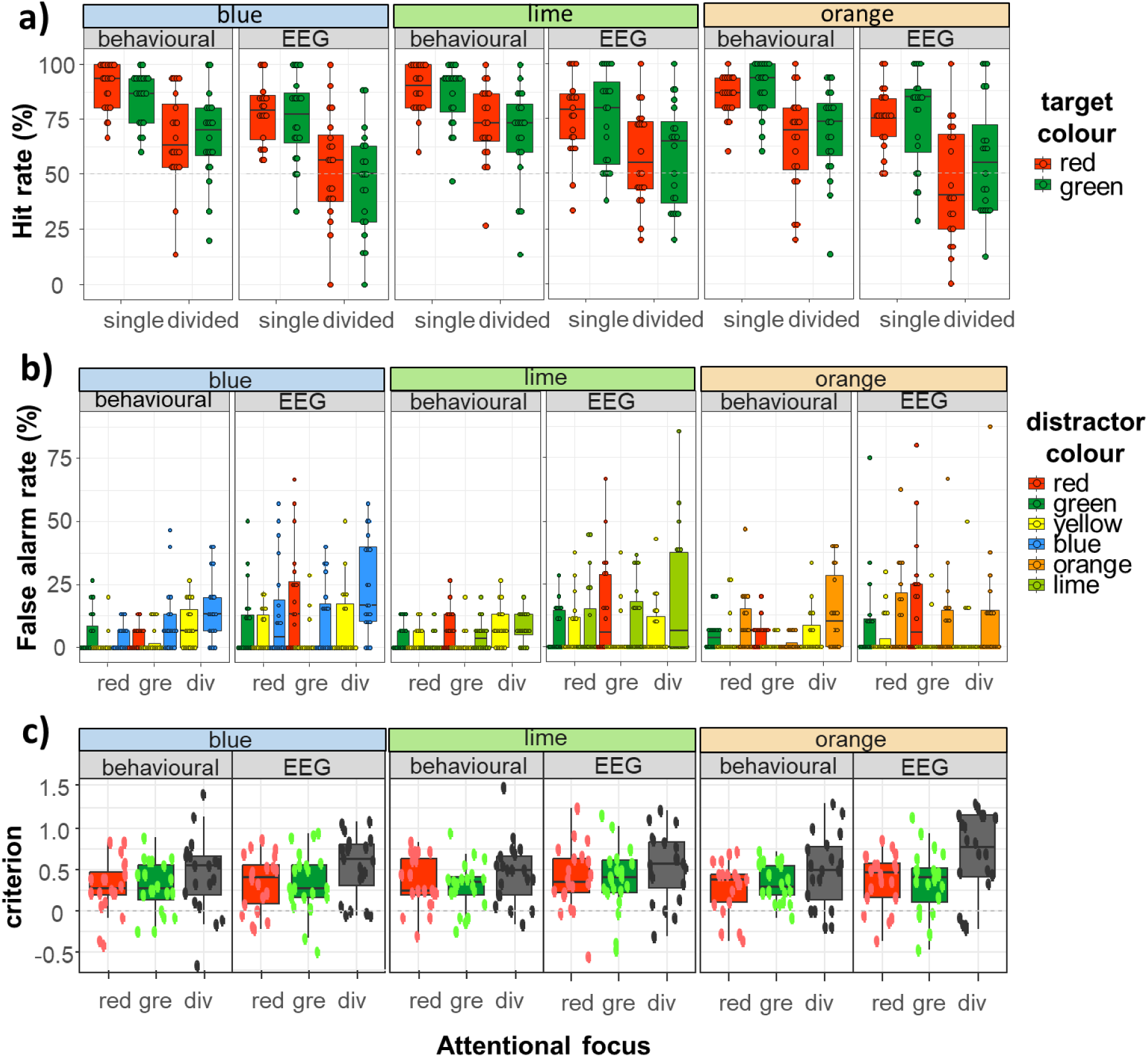
Box plots of behavioural findings: Hit rates (top panel), False Alarm rates (middle panel) and criteria (bottom panel). Within each panel, the data is grouped by experiment (behavioural or EEG) as well as context, determined by the following distractors: blue, orange and lime. Attention could be focused on a single target colour (i.e., red or green, abbreviated as ‘gre’) or divided across these two targets (i.e., divided, abbreviated as ‘div’). To better capture the distribution of the data, we show box plots with individual data points superimposed.

Target-elicited responses were analysed using a GLMM with the following fixed effects - *context (blue, orange or lime), attentional focus (focused on a single target or split between two targets)*, *target colour (red or green)* and *experiment (behavioural or EEG)*. We also added random by-participant intercepts, to account for more basic differences in overall performance. The best-fitting model included attentional focus (χ^2^(1) = 312.39, p < .001), experiment (χ^2^(1) = 113.18, p < .001) and an interaction between target colour and context (χ^2^(2) = 8.964, p=.011; for full details of this model, see Supplementary Materials). As expected, participants were considerably more accurate in detecting target events when they attended a single colour as opposed to when they were asked to divide attention between two targets (OR 3.22, 95% CI 2.81-3.68). The odds of target detection were also substantially increased in the behavioural experiment (OR 2.06, 95% CI 1.8-2.35), indicating interference of stimulus flicker with motion processing. Finally, post-hoc pairwise comparisons of the target colour by context interaction revealed that hit rates for red in the lime context tended to be higher than in the orange context (OR 1.435, 95% CI 1.052 – 1.96, Z = 3.157, p = .013). None of the other pairwise comparisons between contexts reached significance (all ps > .343). To summarise these findings, a major determinant for hit rates appears to be whether attention is focused or divided, with a large cost in performance when attention is split between the two targets. This is consistent with a decreased efficiency of target enhancement. Such a cost could stem from the need to re-allocate central, top-down resources to the monitoring of a secondary attentional template or the continuous need to switch resources between two attentional templates, as they cannot be monitored concurrently. There is also an increase in detection of red targets in the evidently less distracting lime/yellow context compared to the more distracting orange/yellow context, with hit rates being similar across the other contexts.

Responses to yellow distractors were analysed using a GLMM with fixed effects of *context (blue, orange or lime), attentional focus (to red, green or both targets)* and *experiment (behavioural or EEG),* as well as random by-participant intercepts. The best-fitting, final model included an interaction between attentional focus and experiment (χ^2^(2) = 6.325, p = .042), with no contribution of context (see Supplementary materials for detail). Thus, the analysis confirms that the yellow distractor was contextually neutral. We also found some unexpected differences between experiments. Division of attention led to higher false alarms in the behavioural experiment (red vs. divided: OR 0.318, 95% CI 0.163-0.620, Z = - 4.465, p <.001; green vs. divided: OR 0.232, 95% CI 0.110-0.489, Z = −5.309, p <.001), with similar performance across the two foci (red vs. green: OR 1.370, 95% CI 0.574-3.270, Z = 0.982, p = .891). However, in the EEG experiment, division of attention increased the odds of yellow-driven false alarms when compared to the green focus (OR 0.375, 95% CI 0.145-0.968, Z = −2.801, p = .039) but not for red focus (OR 0.813, 95% CI 0.381-1.733, Z = −0.741, p = .965; red and green foci did not statistically differ, OR 2.169, 95% CI 0.833-5.649, Z = 2.192, p = .184). As red and green also acted as distractors when attention had a single focus (i.e., when red was attended, green acted as distractor, and vice versa) we fitted a GLMM with factors of *context* (blue, orange and lime), *distractor colour* (red, green) and *experiment (behavioural or EEG)* as well as random by-participant intercepts to assess any potential differences. As for yellow distractors, we found an interaction between attentional focus and experiment (χ^2^(1) = 6.874, p = .009; see Supplementary Materials for full statistical details), with no differences between the two distractors in the behavioural experiment (OR 1.00, 95% CI 0.583-1.715; Z = 0.00, p = 1.00) but fewer false alarms to green relative to red in the EEG experiment (OR 0.44, 95% CI 0.256-0.755, Z = 3.732, p <.001). This is consistent with lower efficiency of processing red coherent motion events in the EEG experiment and likely to stem from the reduction in motion salience as red flickered at the lowest frequency (8.57 Hz).

For the crucial, context-dependent distractor, there was a significant interaction between all three fixed effects: attentional focus, context and experiment (χ^2^(4) = ]21.377, p<.001; see Supplementary materials). The interaction between focus and context was predicted – we decomposed the statistical effects by comparing these factors within each experiment. In the behavioural experiment, attending to red as opposed to dividing attention between the two targets led to a decrease in blue false alarms (OR 0.198, 95% CI 0.064-0.609, Z = 4.609, p < .001) without robust differences in false alarm rates between focusing on red vs. green (OR 0.314, 95% CI 0.098-1.009, Z = 3.175, p = .051) and dividing attention vs. focusing on green (OR 0.630, 95% CI 0.276-1.435, Z = 1.795, p = .848). Attending red as opposed to dividing attention also led to a decrease in false alarms to lime (OR 0.155, 95% CI 0.027-0.895, Z = −3.401, p = .024) without robust differences in false alarm rates between focusing on red vs. green (OR 0.269, 95% CI 0.043-1.680, Z = −2.293, p = .471) and divided attention vs. green (OR 0.576, 95% CI 0.186-1.786, Z = −1.559, p = .948). Finally, there was a large increase in orange false alarms when attending red compared to green (OR 6.446, 95% CI 1.328-31.29, Z = 3.772, p = .007), while attending green as opposed to dividing attention between the two targets led to a decrease in orange false alarms (OR 0.089, 95% CI 0.019-0.415, Z = 5.023, p < .001; no robust differences for red vs. dividing attention, OR 0.575, 95% CI 0.246-1.341, Z = 2.091, p = .637). Therefore, distraction in divided attention is consistently increased relative to the single focus for the more distal target colour (green target for orange distractor; red target for lime and blue distractors). In addition to that, increased false alarms elicited by the blue distractor when attentional focus included green indicate that in CIELAB, blue is attentionally more proximal to green than to red, invalidating our original assumption that yellow and blue distractors would both provide a supposedly neutral context. On the other hand, there were no robust differences between blue, lime or orange distractors in the EEG experiment (all ps > .075). To understand this discrepancy, it is necessary to look at between-experiment differences. The behavioural experiment had fewer false alarms relative to the EEG experiment but these differences were not uniform. For blue distractors, there were fewer false alarms only for red focus (OR 0.223, 95% CI 0.0634-0.782, Z = −3.825, p = .005, otherwise p>.983) and for orange distractors, fewer false alarms were observed for the green focus (OR 0.129, 95% CI 0.024-0.686, Z = - 3.918, p = .003). Thus, it appears that flicker led to the elevation of false alarm rates in what would have otherwise been a relatively non-challenging condition (compare Fig. 3 and 4), obfuscating previously robust between-condition differences. This further supports our choice to perform both a behavioural and an EEG experiment, with the intention to obtain more valid coherent motion detection data in the absence of flicker.

**Figure 4.**
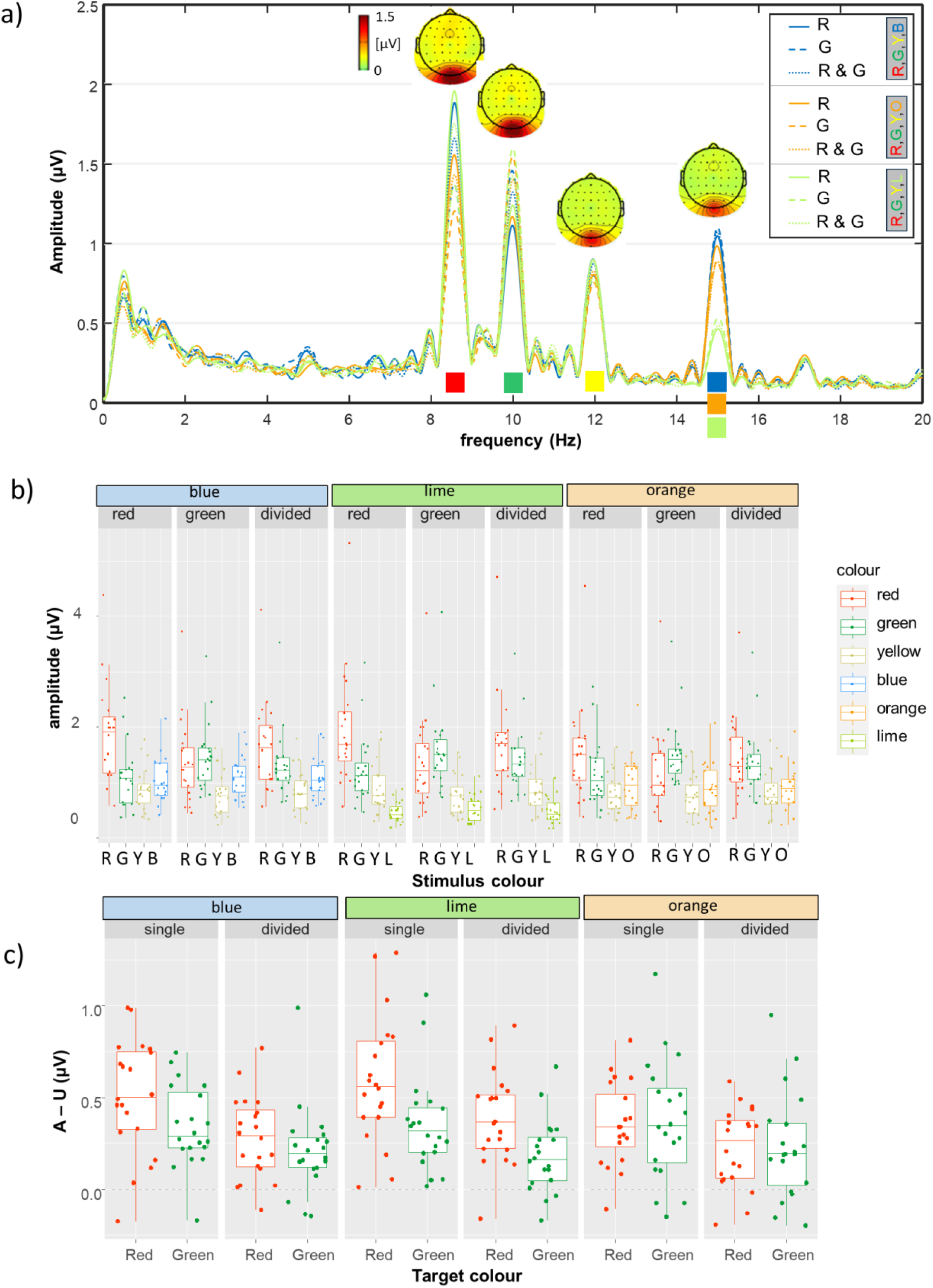
**SSVEP results**. A) Spectra and topographies. Full lines depict amplitudes elicited when red was the target colour, dashed lines depict amplitudes elicited when green was the target colour and dotted lines depict amplitudes when attention was divided between red and green. The colour of the line depicts the context, which in addition to red, green and yellow included either blue, orange or lime. b) Box plot of SSVEP amplitudes, with individual data points overlaid on the graph. c) Box plot of attentional effects calculated from SSVEP amplitudes elicited by green and red targets. Unattended amplitude for the red colour is taken from the ‘attend green’ condition, and vice versa.

Finally, criteria (Fig.3, bottom panel) were fitted with a LMM with fixed effects of *context* (blue, orange and lime), *attentional focus* (red, green, both) and *experiment* (behavioural and EEG), as well as random effect of by-participant intercepts. The best fitting model included fixed effects of *attentional focus* (χ^2^(2) = 32.022, p<.001) and experiment (χ^2^(1)=5.224, p = .022, with EEG experiment being more conservative by 0.072 ± 0.030 SE; for full statistical detail, see Supplementary Materials). A more conservative criterion also manifested itself in the divided attention condition in comparison to attending to red alone (effect size = −0.182 ± 0.037 SE, t(343) = −4.969, p < .001) or green alone (effect size = −0.184 ± 0.037 SE, t(343) = −5.023, p < .001). There was no difference in criteria between the two single foci (effect size = 0.002 ± 0.037 SE, t(343) = 0.054, p = .998). A more conservative criterion for divided attention indicates that the associated increase in false alarms was not due to an adoption of a more liberal response criterion. This is consistent with our observation that divided attention also leads to a large fall in hit rates when compared to focused attention.

In summary, our behavioural data reveals three key patterns: 1) we confirm that flicker selectively decreases the saliency of motion signals for stimuli at lower temporal rates when compared to stimuli at higher temporal rates (Nakayama and Silverman 1984), complicating the comparison of behavioural data for flickering stimuli; 2) we find that divided attention leads to a large reduction of hit rates and a comparatively smaller increase in false alarm rates, as corroborated by the more conservative response criterion; 3) as predicted, false alarms are affected by context, but the assumption that equivalent coordinates in CIELAB space translate to equal attentional selection is not supported. Lime distractors seem less potent in eliciting false alarms than orange distractors, despite being fully matched to them in CIELAB. Additionally, the supposedly neutral blue distractors cause less distraction for red as opposed to green targets.

### EEG experiment: SSVEP results

SSVEP spectra, topographies and amplitudes are depicted in Figure 4a-b. To investigate target enhancement, LMM was fitted to the calculated attentional effects (attended minus unattended SSVEP amplitude; Fig. 4c), with *context (blue, orange or lime), attentional focus (to a single target or divided between two targets)* and *target colour from which responses are analysed (red or green)* as fixed effects and participants as random effects (see Supplementary materials for full details of the best fitting model). *Attentional focus* had the biggest influence on SSVEP attentional modulation (χ^2^(1) = 31.318, p < .001), with an average loss of about 50% (i.e. 0.169 µV) when dividing attention. *Context* and *target colour* interacted (χ^2^(2) = 11.005, p = .004), qualifying the main effects observed for each of these factors individually. Further examination of this interaction revealed that the attentional effect for red in the lime context was higher when compared to red in the orange context (estimate = 0.202 µV, t(226) = 3.956, p = .001) but not the blue context (estimate = 0.096 µV, t(226) = 1.881, p = .417); orange and blue contexts did not differ robustly either (estimate = 0.106 µV, t(226) = 2.075, p = .3044). Meanwhile, attentional effects for green remained relatively stable, not robustly affected by any of the contexts (all ts < 0.718, ps > .980). Thus, higher target enhancement for red was possible in the presence of yellow and lime. We also performed the statistical analysis separately on red and green data, to avoid any confounds due to comparing data stemming from two stimulation frequencies within the same model. This analysis confirmed all the effects reported above (i.e., for red, there were main effects of both context and focus, while green amplitude was only affected by focus; see Supplementary Materials for statistical tables).

To investigate how distractors were processed between contexts, we first focused on the SSVEP amplitude elicited by yellow, which was the distractor that was constantly present (see Figure 4b). A LMM with *context (blue, orange or lime),* and *attentional focus (to red, green or divided)* as main effects and participants as random effects was fitted (full statistical details are in the Supplementary Materials). The best fitting model contained the additive effects of *context* and *attentional focus*. Amplitude for yellow was lower in the orange context (p = .003), with similar amplitudes across blue and lime contexts (p = .928). In addition, amplitude was lower when attention was focused on green, compared to when attention was focused on red (p <.001; red vs. divided attention not significantly different, p = .987).

Note that in Fig. 4a-b amplitudes elicited by blue, orange and lime at 15 Hz are markedly different. Therefore, we fitted LME models with *attentional focus (red, green or divided)* as a fixed effect and participants as random effects for each of the changeable distractor colours separately, intending to reveal whether their neural processing was affected by the target colour(s). For orange, attentional focus contributed significantly to the model (χ^2^(2) = 5.587, p = .007), with higher amplitudes when attending red compared to attending green (t(42.1) = 2.821, p = .0195) or dividing attention between red and green (t(42.1) = 2.822, p = .0194; green vs. divided attention, t(42.1) = 0.001, p = 1.00). For blue and lime, attentional focus did not play a significant role in explaining any of the variance.

In a final analysis, we evaluated if target-related attentional increases at the neural level follow a simple resource allocation model. If this were the case, attentional modulation in divided attention conditions should be half of its magnitude when only single colours are attended (Adamian *et al*. 2019; Adamian and Andersen 2022; Andersen, Müller & Hillyard, 2009). Alternatively, if these were implemented independently for each colour, akin to selection of feature values from different dimensions (Andersen *et al*. 2015; Adamian *et al*. 2019), attentional modulation in the divided attention condition would be as large as in the attend single colour conditions. Predictions based on simple resource allocation (i.e., 50% of attentional increase when dividing attention) seem to fit the data extremely well, only slightly underestimating the actual attentional modulation of SSVEP amplitudes. The multiplier *p*, reflecting the reduction in attentional modulation between single and divided attentional foci ( AmpDiv_colour_ = p * (AmpColour_red_ + AmpColour_green_), ranged between 0.511 and 0.538, indicating that attentional resources are approximately halved between the two colours (Red, blue context: 0.516, 95% CI 0.486 – 0.545, t(19) = 1.125, p = .275; Green, blue context: 0.522, 95% CI 0.492 – 0.552, t(19) = 1.563, p = .135; Red, orange context: 0.536, 95% CI 0.500 – 0.572, t(19) = 2.099, p = .0494; Green, orange context: 0.520, 95% CI 0.477 – 0.564, t(19) = 0.983, p = .338; Red, lime context: 0.538, 95% CI 0.514 – 0.562, t(19) = 3.35, p = .003; Green, lime context: 0.511, 95% CI 0.483 – 0.540, t(19) = 0.843, p = .41). Summation coefficient for red targets in orange and lime contexts is statistically significantly above 0.5, but this deviation is very small (∼4%).

## Discussion

We investigated whether concurrent selection of two colour targets can be conceived as a process in which two attentional foci are applied within the multidimensional colour space, enacting control on the distribution of processing resources across colour representations on the basis of their proximity within this space. To achieve this, we designed an experiment that manipulated both the number of colour targets (one or two) and the proximity of distractors to the targets in a perceptual colour space (intermediate or closer to one of the targets). In a sustained multi-coloured RDK coherent motion detection task, we placed red and green targets in one of three contexts: i) a neutral context with intermediate distractors, falling between the two target hues in CIELAB space (blue and yellow); ii) a context with one distractor closer to red (orange and yellow); iii) a context with one distractor closer to green (lime and yellow). In the behavioural experiment, we found that hit rates were overwhelmingly dependent on whether attention was focused or divided, confirming that concurrent selection of two colours was possible, but came at a significant cost to the efficiency of target enhancement. On the other hand, false alarms were more strongly affected by context: blue, orange and lime distractors affected the performance differentially. In line with this, target-driven SSVEP data were consistent with a combination of two separate foci that controlled the distribution of attention across elements in the scene, with allocated resources being approximately halved with divided attention. On the other hand, representational proximity between targets and distractors affected distractor-elicited neural responses to a much lesser degree compared to behavioural responses. This might indicate that concurrent selection of two categorically distinct colours is mainly limited by a central, resource-related bottleneck, with more minor contributions from a local, perceptual-distance-related bottleneck.

As predicted, we found higher rates of responses to distractors in contexts that put them closer to one colour, e.g. orange elicits more false alarms when the target is red and lime elicits more false alarms when the target is green. However, contrary to our predictions, this did not occur evenly across our contexts, which were presumed to be either neutral (blue), closer to red (orange) or closer to green (lime). Blue was found to elicit more false alarms when the target was green, while the overall number of false alarms was the lowest (i.e., most ‘neutral’) in the yellow/lime context. This unexpected finding implies that while CIELAB may be perceptually uniform, this does not guarantee in any way its uniformity in terms of attentional effects or categorical representativeness of different hues. Unlike simple colour discrimination (for a discussion, see Martinovic et al. 2020), performance in a visual search task for colour targets is influenced by categorical status (Found and Muller 1996; for an overview, see Martinovic and Hardman 2016). Categories can also be used to set attentional guidance based on an element’s position in relation to the other colours in the display (e.g. ‘redder than’ or ‘yellower than’, Becker 2010; such relational colour selection is likely to be an aspect of good-enough attentional guidance, Yu *et al*. 2023). We observed higher levels of distractor interference, both behaviourally and neurally, for the orange colour as compared to the lime colour. This asymmetry is not consistent with positions of our stimulus colours in CIELAB nor with their positions in relation to unique hues (see Fig. 1), but is consistent with categorical labelling data, which indicates a larger overlap between orange and red (Malkoc et al. 2005, Fig. 1). The fact that CIELAB may be perceptually but not attentionally uniform poses problems for studies that probe multiple locations in CIELAB but subsequently collapse the data across hues (e.g. Stormer and Alvarez 2014).

In a sustained task such as ours, proximity to categorical best-exemplars may act directly, by facilitating the efficient setting and maintenance of attentional templates (Bae et al. 2015), or indirectly, by facilitating grouping of more similar hues together, be they targets or distractors (Matera et al. 2020). Since the orange distractor is more similar to red than to yellow, this may lead to a spread of activation in accordance with the feature similarity gain model (Martinez-Trujillo and Treue 2004; Oxner et al. 2023), making it more difficult to allocate top-down resources (i.e. to successfully bias competition) towards the red target whilst ignoring orange. On the other hand, if the ‘yellowness’ of the lime distractor can be successfully used to group it together with yellow and away from green, this would make it easier to bias competition towards the green target. Grouping of local motion signals by colour appears to precede global motion processing and is under attentional control (Martinovic et al. 2009; Wuerger et al. 2011). If colour similarity increases the grouping of colours in the RDK, this would create predicted costs when they need to be selected or ignored independently (e.g., red/orange) but also produce unexpected benefits when they need to be ignored together (e.g., yellow/lime). In fact, the special status of colour as an extremely salient feature dimension in terms of attention (see Found and Muller 1996) is at least partly derived by our ability to maintain and use multiple templates in short-term memory that are based on a colour’s categorical status, mediated through ease of verbal labelling (Forsberg et al. 2019). As discussed in a recent review, this multidimensionality of colour is likely to have important repercussions for colour-based attentional selection (Liesefeld and Müller 2019).

In a recent combined SSVEP-fMRI study, attentional colour modulations were mainly observed in area V4v (Boylan et al. 2023). Together with VO1 and VO2, this is the area in which colour selection has maximal effects (Thayer and Sprague 2023). In V4, colour representations are organised in a way that follows the perceptual hue circle but also exhibits categorical clustering (Brouwer and Heeger 2009, 2013). A model has been proposed that attributes this clustering to a colour-specific gain change: the gain of each neuron changes as a function of its selectivity relative to the centres of the colour categories. If attentional selection in extra-striate cortices is implemented through stimulus-stimulus interactions (for a discussion of this model, see Parker 2020), then attentional modulations would be more effective if aligned with category centres, both in terms of target enhancement and distractor suppression. This is a testable prediction which can be directly examined in future studies, manipulating category representativeness of targets and distractors systematically.

Returning to the highly congruent effects observed in behavioural and neural data elicited by targets, the ∼20-25% cost in hit rates for divided attention corresponded to a ∼50% reduction in amplitude that was relatively well predicted by a simple averaging of the amplitudes elicited by red and green in the two focused attention conditions. Processing is altered not only for targets, but also for some distractors: orange, which elicited highest behavioural interference when red was attended, also had higher amplitude if the focus of attention was on red alone. However, the behavioural costs were comparatively smaller than for targets, and neural changes were even more limited - 9% change in amplitude for orange distractors and no discernible neural changes for lime or blue distractors, despite increased false alarms. While attentional modulations of sensory coding and behavioural readout are both likely to have some neural correlates in V4 (Parker 2020), behavioural readout would also be influenced by multiple further components of processing beyond mere competitive interactions in visual cortices (e.g., the dorsal frontoparietal network, Liu 2019). This implies that the need to maintain two active target templates poses higher costs at later stages of attentional processing. Finally, we observe that orange-driven amplitude is similarly reduced when attentional focus is on green or divided between red and green. Thus, it appears that having a green target template is sufficient to eliminate the allocation of visuo-cortical resources to orange that stems from the adoption of a red target template. In the context of limited parallel or limited serial models of multiple-target selection (Ort and Olivers 2020), serial switching between the two templates during selection would mean that the spreading of perceptual resources to orange should be halved, rather than eliminated. Future studies should attempt to precisely quantify the residual resources allocated to orange whilst attending to red, to provide more decisive evidence disambiguating between parallel (already favoured in the literature) or serial selection models.

Putting these findings together, we have shown that target enhancement in divided attention to colour can be predicted from single colour selection. Colour distances in CIELAB space only partially account for our results, both in terms of behavioural and neural data. The findings of our study pave the way for a more refined exploration of the relations between categorical status, colour content (e.g., as identified by hue scaling) and perceptual differences. Should full understanding of these relations be obtained, it may be possible to quantify an attentional colour space, which could be used to predict the ease with which colours can be selected together or filtered out as distractors.

## Open science

Raw behavioural and averaged EEG data, as well as statistical analysis scripts in R are publicly available on Open Science Framework (OSF): https://osf.io/2kjsx/

## Acknowledgments

AB was supported by a BBSRC Eastbio research experience placement (Doctoral Training Programme BB/M010996/1) to work in the lab of JM. The authors state no conflict of interest. For the purpose of open access, the authors have applied a Creative Commons Attribution (CC BY) licence to any Author Accepted Manuscript version arising from this submission.

## Author contributions

JM designed the experiment; JM and SKA implemented the experimental and analysis scripts; AB and JM collected the data; JM analysed the data; AB, JM and SKA wrote the original version of the manuscript; JM and SKA revised the manuscript.

## Supplementary Materials

### Supplementary Material – Best fitting statistical models

The tables present estimated parameters, their standard errors or confidence intervals, z-values, p-values and effect sizes for the best fitting GLMM or LMM.

#### 1 Behavioural results

**Table S1.**
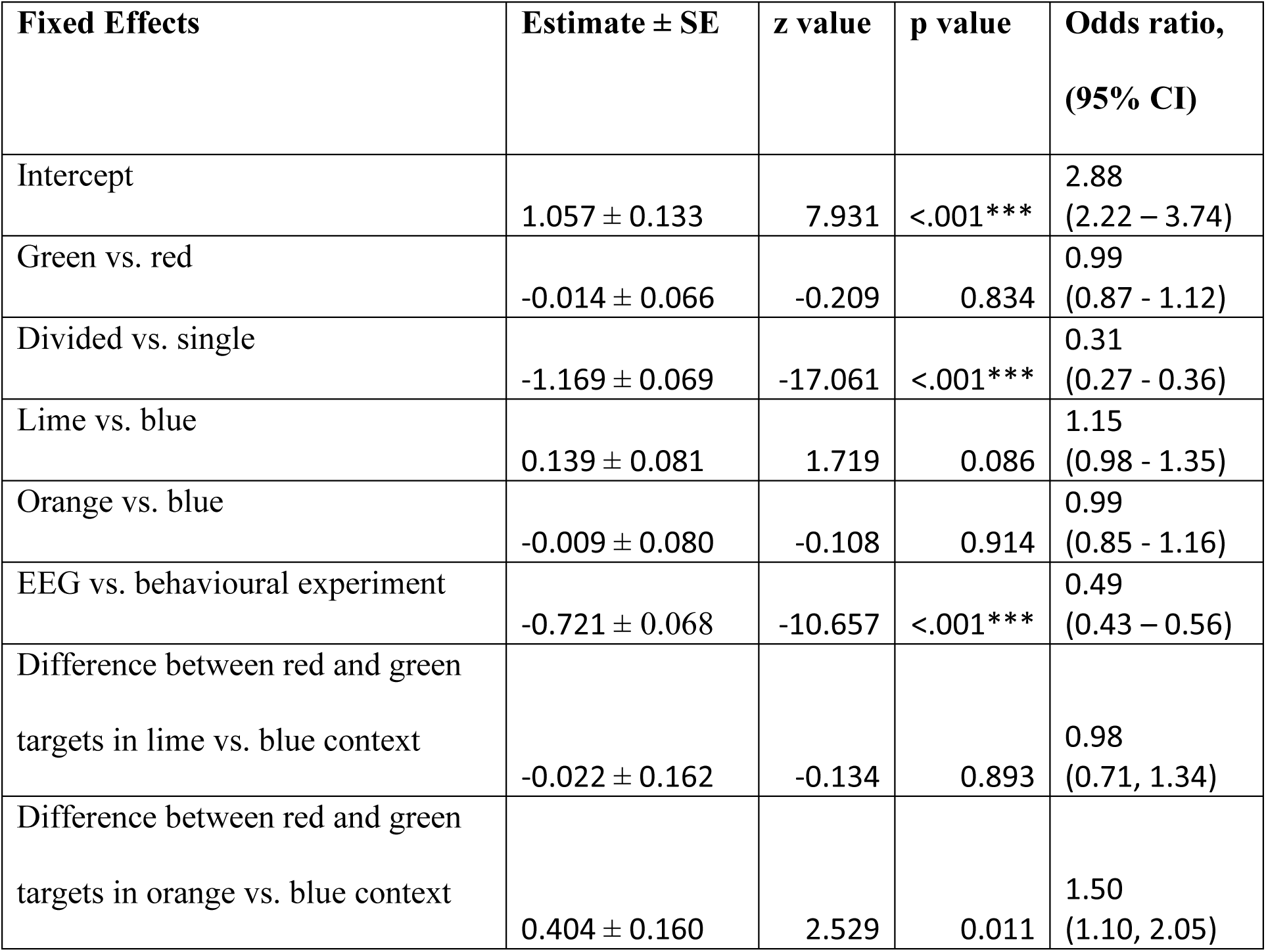
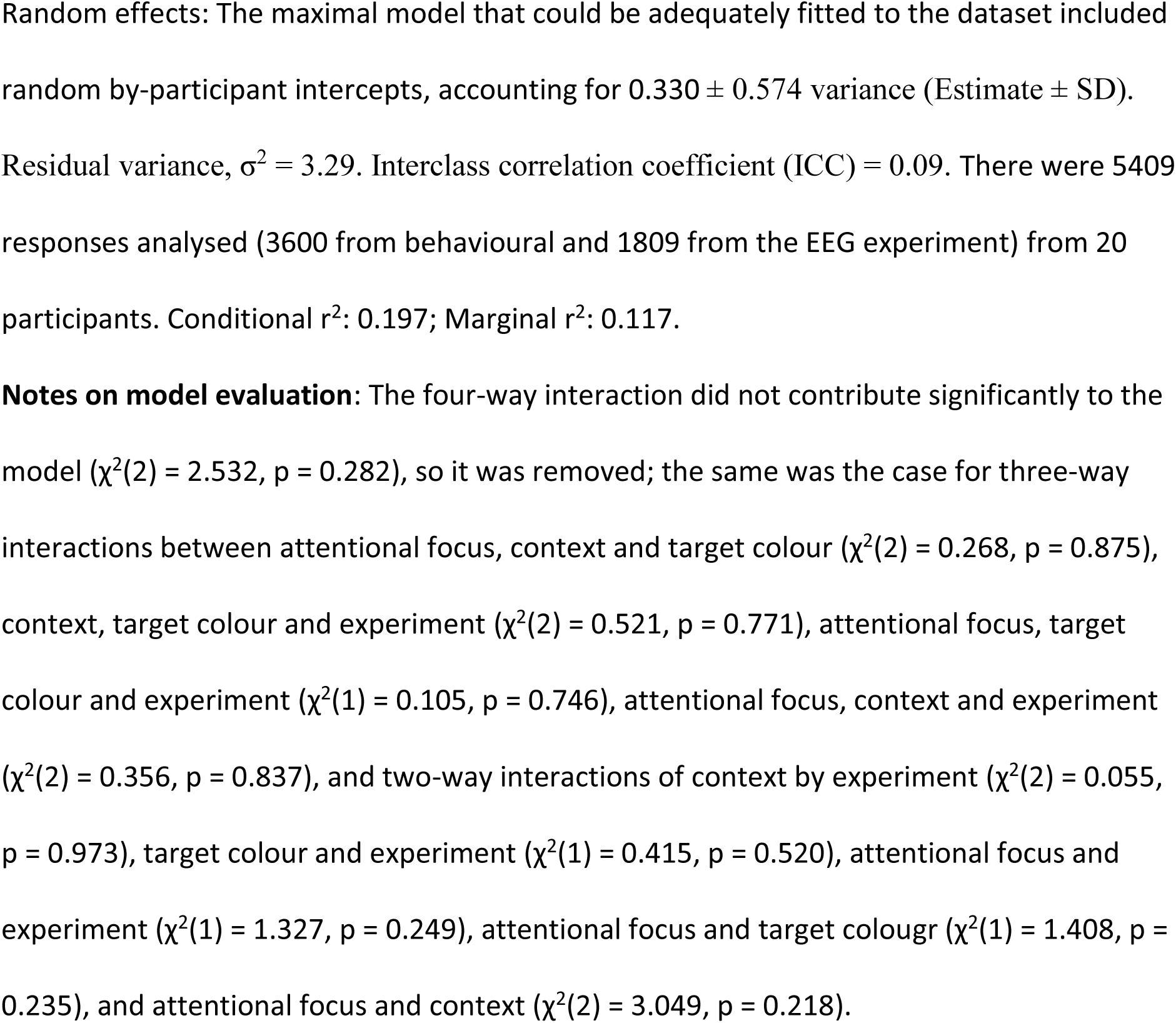
GLMM - Responses to target events. Reference levels were set to single focus, red target, blue context and behavioural experiment. Model equation: response ∼ 1 + target_colour + attentional_focus + colour_context + experiment + target_colour : colour_context + (1|participant)

**Table S2.** GLMM - Responses to yellow distractors. Reference levels were set to red focus and behavioural experiment. Model equation: response ∼ 1 + target_colour + attentional_focus + experiment + target_colour : experiment + (1|participant)

| Fixed Effects | Estimate ± SE | z value | p value | Odds ratio, (95% CI) |
| --- | --- | --- | --- | --- |
| Intercept | -3.551 ± 0.234 | -15.185 | <.001*** | 0.03<br>(0.02 - 0.05) |
| Green vs. red | -0.545 ± 0.239 | -2.282 | 0.022 | 0.58<br>(0.36 - 0.93) |
| Divided vs. red | 0.677 ± 0.187 | 3.629 | <.001*** | 1.97<br>(1.37 - 2.84) |
| EEG vs. behavioural experiment | 0.382 ± 0.177 | 2.158 | 0.031 | 1.47<br>(1.04 – 2.07) |
| Green vs. red / EEG vs.<br>behavioural experiment | -0.459 ± 0.477 | -0.962 | 0.336 | 0.63<br>(0.25 – 1.61) |
| Divided vs. red / EEG vs.<br>behavioural experiment | -0.940 ± 0.373 | -2.522 | 0.012 | 0.39<br>(0.19 – 0.81) |

Random effects: The maximal model that could be adequately fitted to the dataset included random by-participant intercepts, which accounted for $0.829 \pm 0.911$ in variance (Estimate ± SD). Residual variance 3.29 (ICC = 0.20). There were 4042 responses analysed (2700 from the behavioural and 1342 from the EEG experiment), from a total of 20 participants. Conditional $r^2$ : 0.265; Marginal $r^2$ : 0.079.
**Note:** The three-way interaction did not contribute significantly to the model ( $\chi^2(2) = 2.5318$ , $p = 0.282$ ), so it was removed; the same was the case for two-way interactions between attentional focus and context ( $\chi^2(4) = 4.415$ , $p = 0.353$ ), context and experiment ( $\chi^2(2) = 0.259$ , $p = 0.879$ ), and the fixed effect of context ( $\chi^2(2) = 2.384$ , $p = 0.304$ ).

**Table S3.** GLMM - Responses to red and green distractors. Reference levels were set to red colour and behavioural experiment. Model equation: response ∼ 1 + target_colour + experiment + target_colour : experiment + (1|participant)

| Fixed Effects | Estimate ± SE | z value | p value | Odds ratio,<br>(95% CI) |
| --- | --- | --- | --- | --- |
| Intercept | -2.787 ± 0.219 | -12.76 | <.001*** | 0.06<br>(0.04, 0.09) |
| Green vs. red colour | 0.411 ± 0.156 | 2.641 | 0.008 | 1.51<br>(1.11-2.05) |
| EEG vs. behavioural experiment | 1.030 ± 0.156 | 6.593 | <.001*** | 2.80<br>(2.06-3.80) |
| Green vs. red across EEG vs.<br>behavioural experiment | 0.822 ± 0.311 | 2.641 | 0.008 | 2.27<br>(1.24 – 4.19) |

Random effects: The maximal model that could be adequately fitted to the dataset included random by-participant intercepts, which accounted for $0.763 \pm 0.873$ in variance (Estimate ± SD). Residual variance 3.29 (ICC = 0.19). There were 2689 responses analysed, from a total of 20 participants. Conditional $r^2$ : 0.243; Marginal $r^2$ : 0.067.
**Note:** The three-way interaction did not contribute significantly to the model ( $\chi^2(2) = 1.415$ , $p = 0.493$ ), so it was removed; the same was the case for attentional focus by context ( $\chi^2(2) = 3.497$ , $p = 0.174$ ) context by experiment ( $\chi^2(2) = 2.152$ , $p = 0.341$ ) and context ( $\chi^2(2) = 0.148$ , $p = 0.929$ ).

**Table S4.**
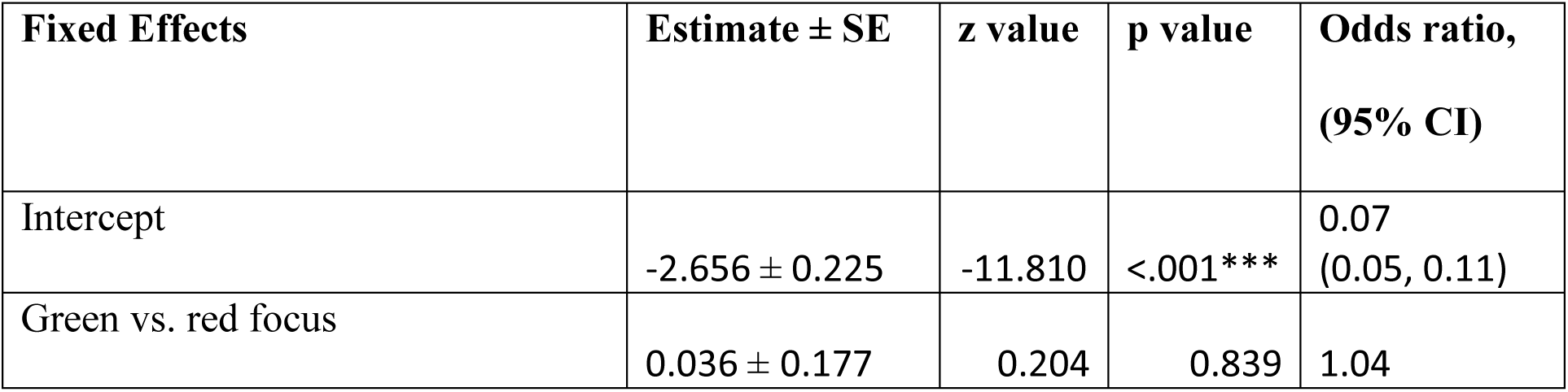
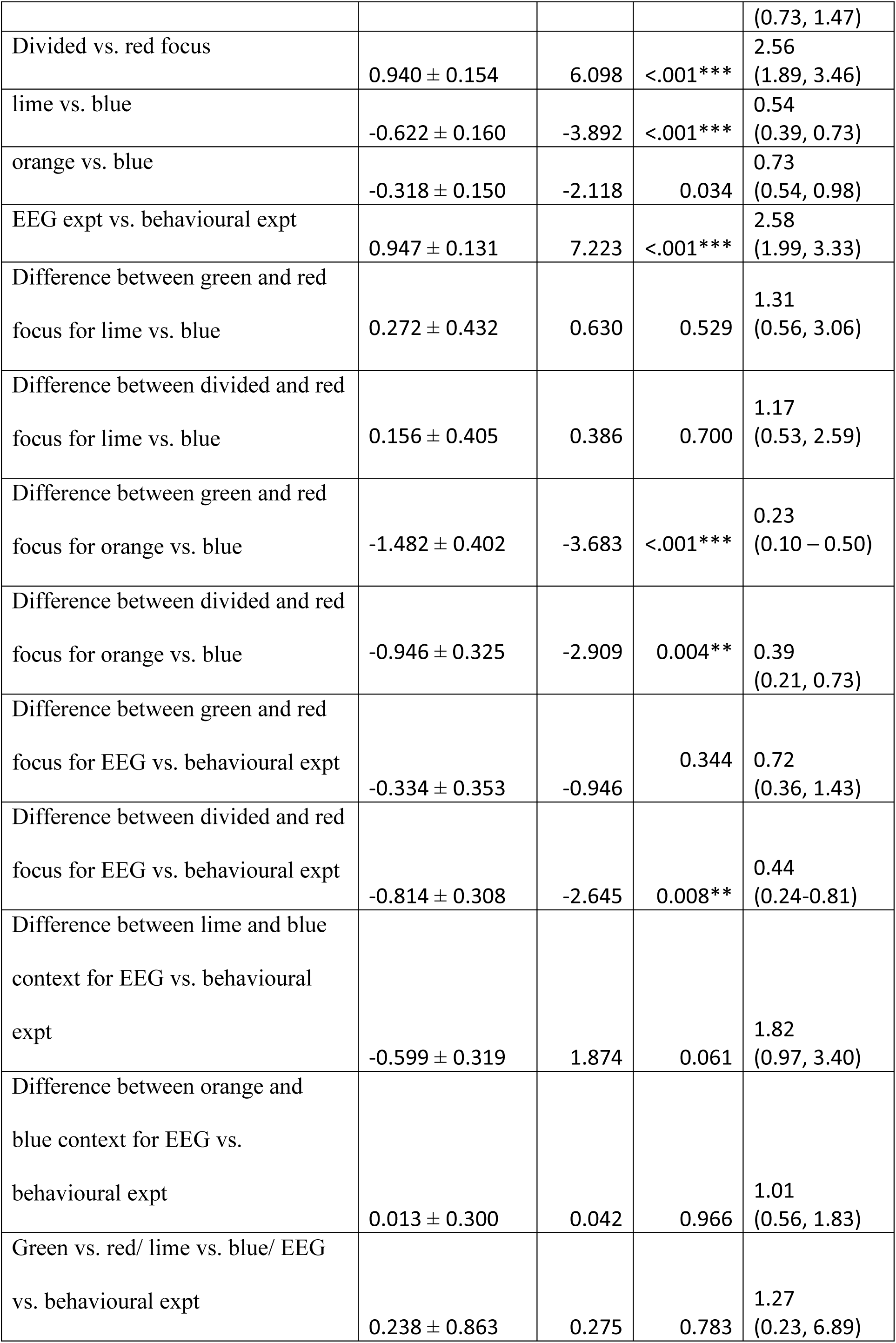
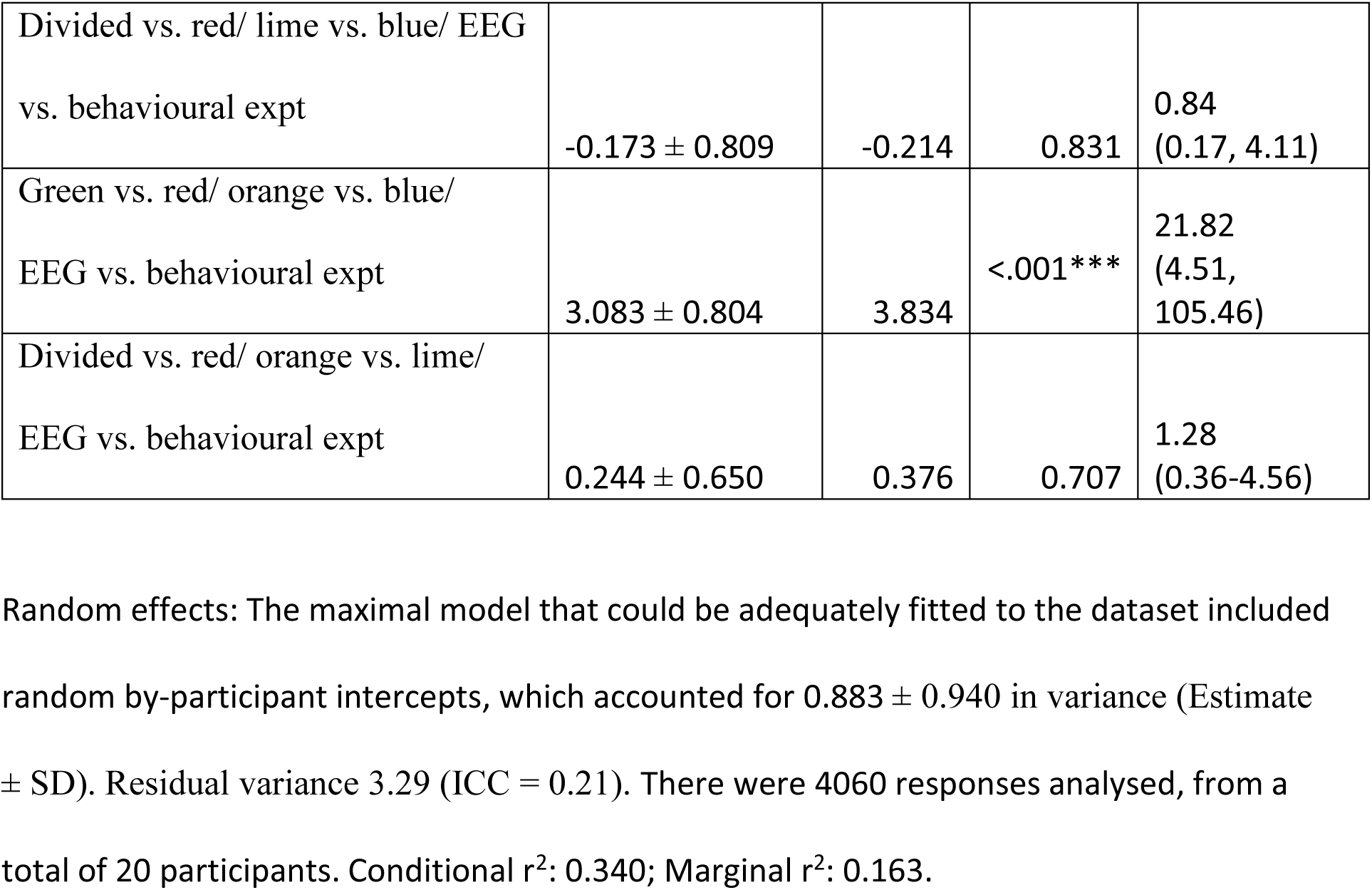
Responses to blue, orange and lime distractors. Reference levels were set to red focus, blue distractor and behavioural experiment. Model equation: response ∼ 1 + attentional_focus + colour_context + experiment + attentional_focus:colour_context + attentional_focus:experiment + attentional_focus:colour_context:experiment + (1|participant)

**Table S5.**
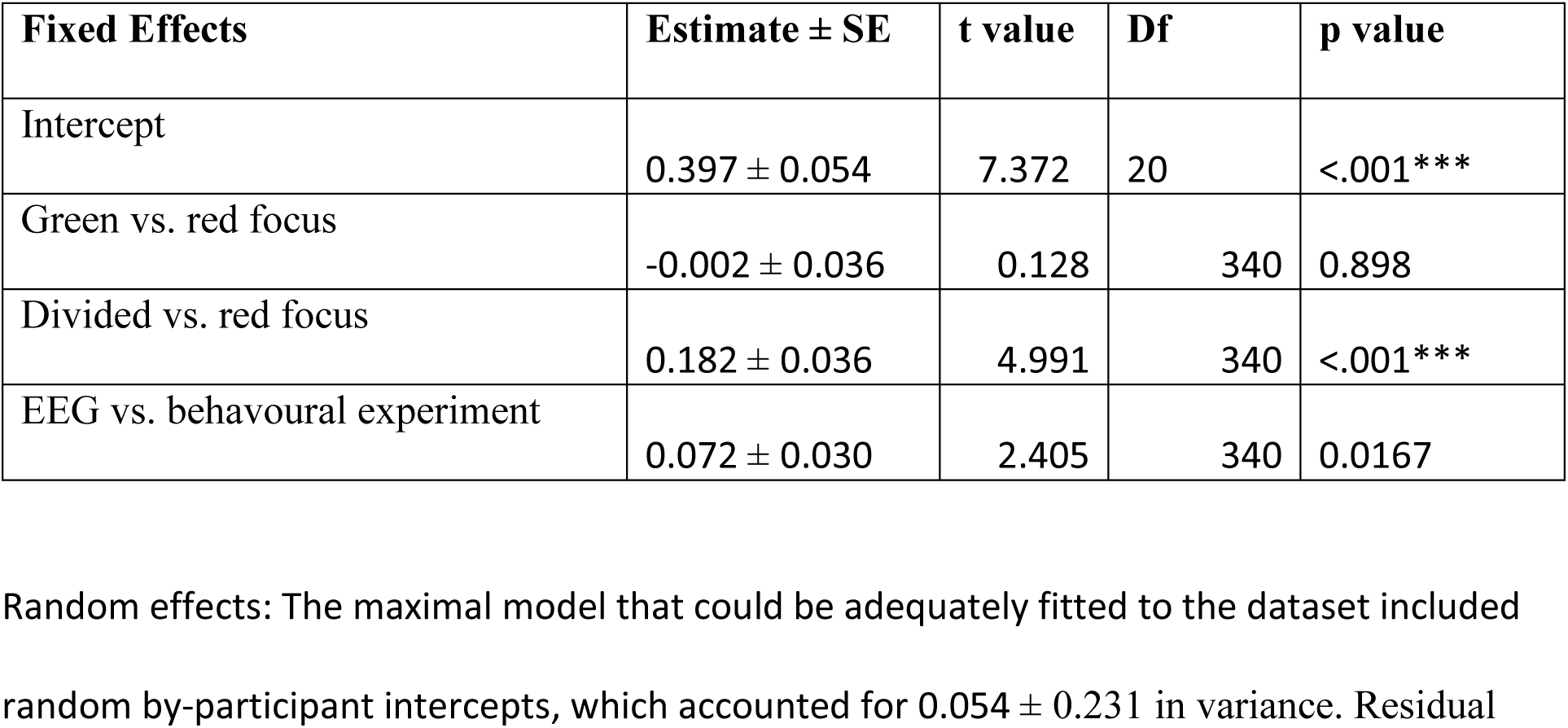
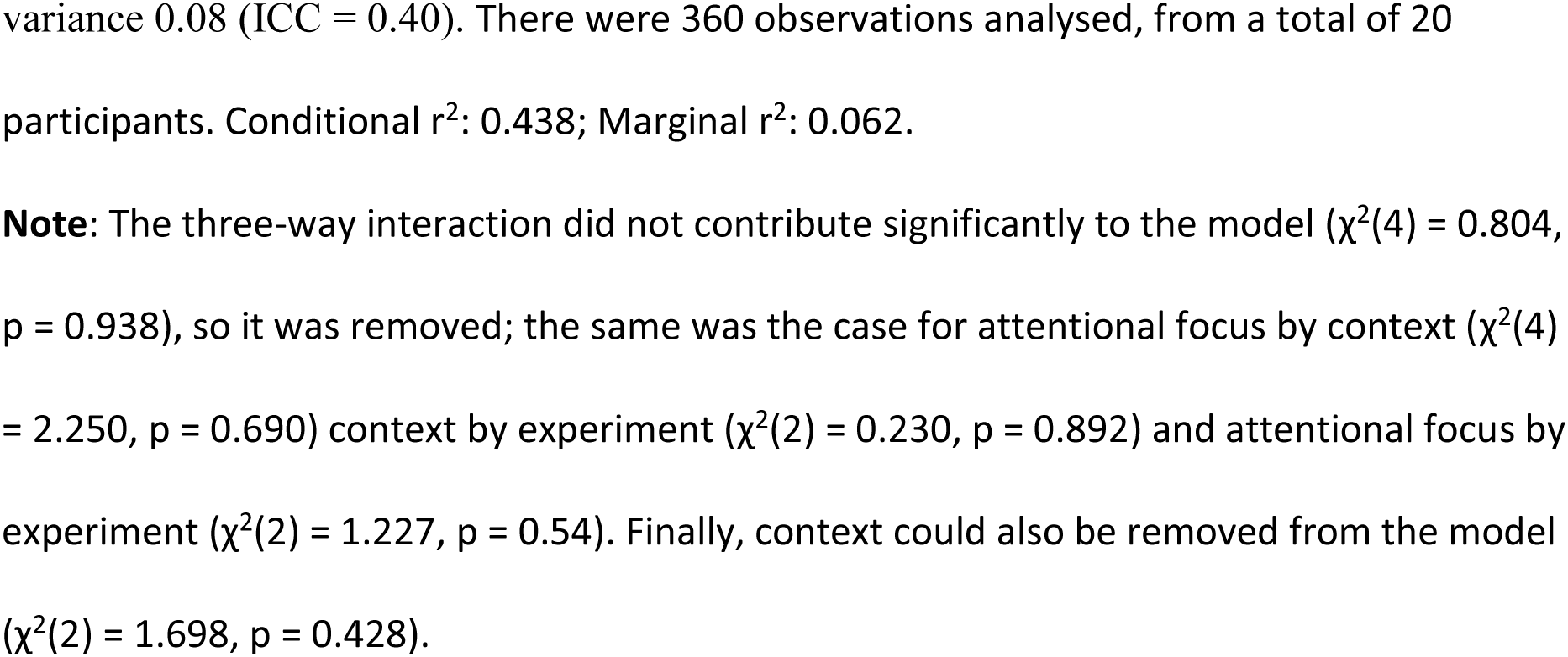
LMM - Response criteria. Reference levels were set to red focus and blue context. Model equation: C ∼ 1 + attentional_focus + experiment + (1|participant)

**Table S6.** *LMM* - attentional effects on SSVEP amplitudes. Reference levels were set to red focus and blue context. Model equation: attended minus unattended target amplitude ∼ 1 + attentional_focus + target_colour + colour_context +target_colour:colour_context + (1|participant)

|  | Attentional modulation (A-U) |  |  |  |
| --- | --- | --- | --- | --- |
| *Predictors* | *Estimate* | *95% CI* | *t* | *p* |
| (Intercept) | 0.338 | 0.28 – 0.40 | 10.752 | <.001 |
| Attentional Focus (divided vs. single) | -0.169 | -0.23 - -<br>0.11 | -5.801 | <.001 |
| Target Colour (green vs. red) | -0.109 | -0.17 - -<br>0.05 | -3.757 | <.001 |
| Context (lime vs. blue) | 0.044 | -0.03 – 0.11 | 1.226 | 0.221 |
| Context (orange vs. blue) | -0.039 | -0.11 – 0.03 | -1.095 | 0.274 |
| Target Colour x Lime vs. blue context difference | -0.105 | -0.24 – 0.04 | -1.471 | 0.142 |
| Target Colour x Orange vs. blue context difference | 0.134 | -0.01 - 0.27 | 1.880 | 0.061 |
| *Random effects* |  |  |  |  |
| Residual variance, $\sigma^2 = 0.051$ | | | | |
| Participant variance 0.0156 (0.125 SD) |  |  |  |  |
| ICC 0.24 |  |  |  |  |
| N participant 20, 240 Observations in total |  |  |  |  |
| Marginal $R^2$ / Conditional $R^2$ 0.171 / 0.366 | | | | |

**Note:** The three-way interaction did not contribute significantly to the model ( $\chi^2(2) = 0.565$ , $p = 0.754$ ), so it was removed; the same was the case for attentional focus by context ( $\chi^2(2) = 0.581$ , $p = 0.748$ ) and attentional focus by colour ( $\chi^2(1) = 0.396$ , $p = 0.529$ ).

**Table S6.1.** *LMM -* attentional effects on SSVEP amplitudes – separate analysis for red. Reference level set to blue context. Model equation: attended minus unattended red amplitude ∼ 1 + attentional_focus + colour_context + (1|participant)

|  | Attentional modulation (A-U) |  |  |  |
| --- | --- | --- | --- | --- |
| *Predictors* | *Estimate* | *95% CI* | *t* | *p* |
| (Intercept) | 0.393 | 0.31 – 0.48 | 9.377 | <.001 |
| Attentional Focus (divided vs. single) | -0.187 | -0.26 - -<br>0.11 | -5.135 | <.001 |
| Context (lime vs. blue) | 0.096 | 0.01 – 0.18 | 2.154 | 0.033 |
| Context (orange vs. blue) | -0.106 | -0.19 – -<br>0.02 | -2.375 | 0.019 |
| *Random effects* |  |  |  |  |
| Residual variance, $\sigma^2 = 0.040$ | | | | |
| Participant variance 0.028 |  |  |  |  |
| ICC 0.42 |  |  |  |  |
| N participant 20, 120 Observations in total |  |  |  |  |
| Marginal $R^2$ / Conditional $R^2$ 0.187 / 0.526 | | | | |

**Note:** The two-way interaction did not contribute significantly to the model ( $\chi^2(2) = 1.257$ , $p = 0.534$ ), so it was removed.

**Table S6.2.**
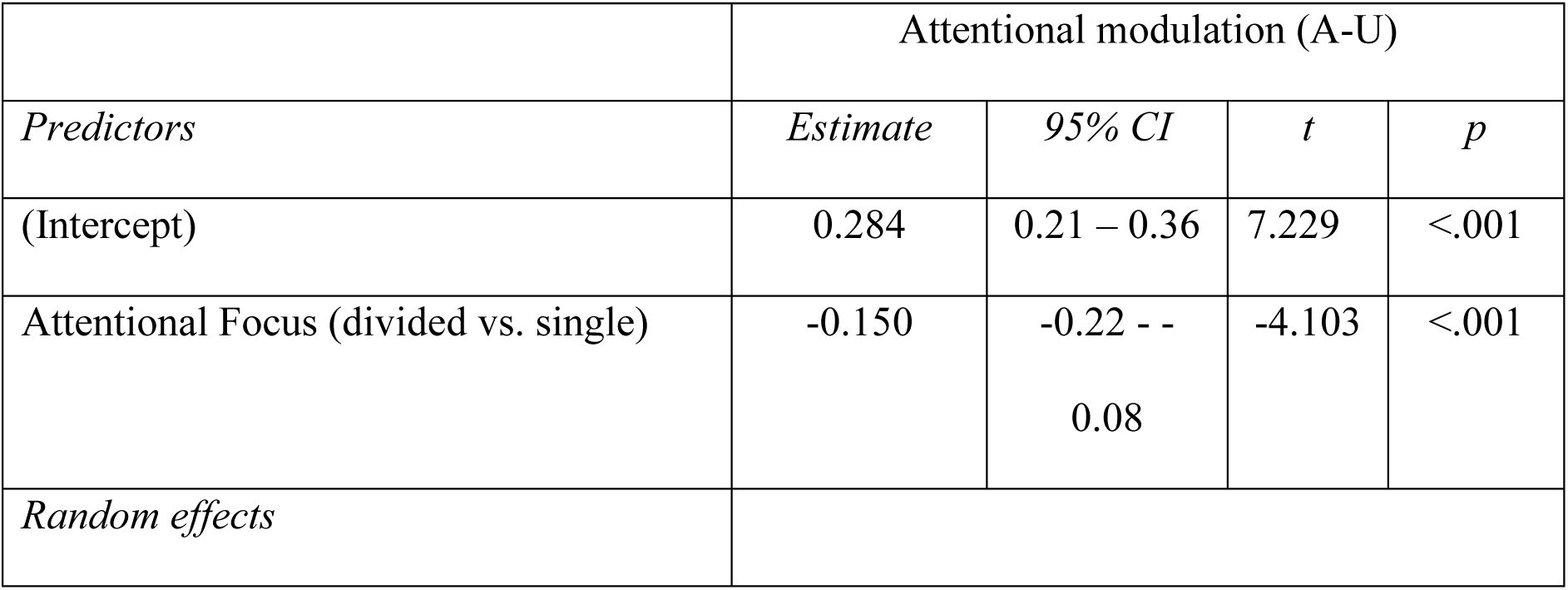
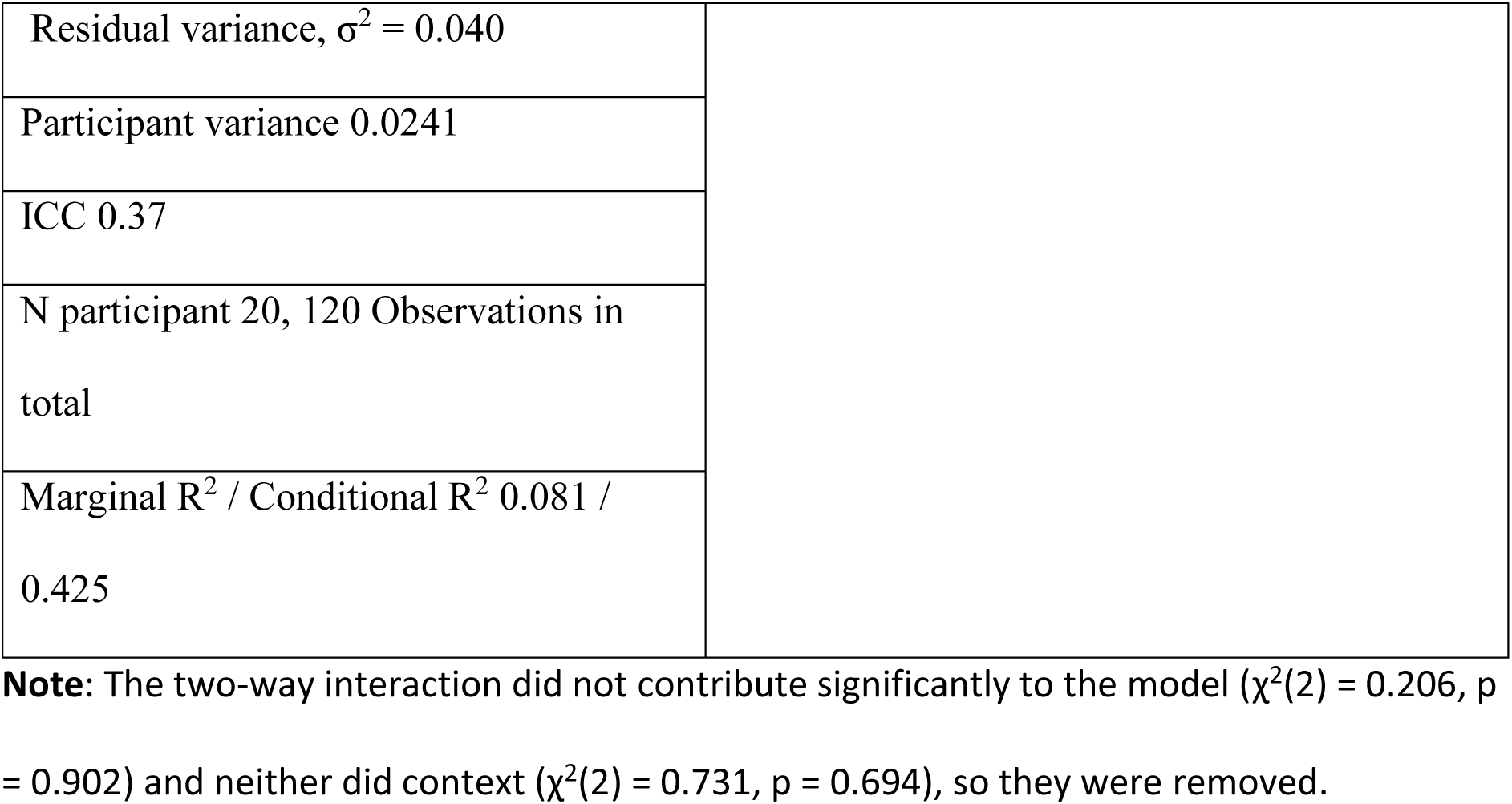
*LMM -* attentional effects on SSVEP amplitudes – separate analysis for green. Reference level set to single focus. Model equation: attended minus unattended green amplitude ∼ 1 + attentional_focus + (1|participant)

|  | Attentional modulation (A-U) |  |  |  |
| --- | --- | --- | --- | --- |
| *Predictors* | *Estimate* | *95% CI* | *t* | *p* |
| (Intercept) | 0.284 | 0.21 – 0.36 | 7.229 | <.001 |
| Attentional Focus (divided vs. single) | -0.150 | -0.22 - -<br>0.08 | -4.103 | <.001 |
| *Random effects* |  |  |  |  |

|  |  |
| --- | --- |
| Residual variance, $\sigma^2 = 0.040$ | |
| Participant variance 0.0241 |  |
| ICC 0.37 |  |
| N participant 20, 120 Observations in total |  |
| Marginal $R^2$ / Conditional $R^2$ 0.081 / 0.425 | |

**Note:** The two-way interaction did not contribute significantly to the model ( $\chi^2(2) = 0.206$ , $p = 0.902$ ) and neither did context ( $\chi^2(2) = 0.731$ , $p = 0.694$ ), so they were removed.

**Table S7.** *LMM -* SSVEP amplitudes elicited by yellow. Reference levels were set to red focus and blue context. Model equation: amplitude ∼ 1 + attentional_focus + colour_context + (1|participant)

|  | Amplitude |  |  |  |
| --- | --- | --- | --- | --- |
| *Predictors* | *Estimate* | *95% CI* | *t* | *p* |
| (Intercept) | 0.831 | 0.704 –<br>0.964 | -2.904 | <.001 |
| Attentional Focus: green vs. red | -0.083 | -0.115 - -<br>0.052 | -4.361 | <.001 |
| Attentional Focus: divided vs. red | 0.0003 | -0.035 -<br>0.036 | -0.423 | 0.987 |
| Context (lime vs. blue) | 0.002 | -0.037 –<br>0.040 | -0.293 | 0.928 |
| Context (orange vs. blue) | -0.057 | -0.102– -<br>0.013 | -2.927 | 0.003 |
| *Random effects* |  |  |  |  |
| Residual variance, $\sigma^2 = 0.0228$ | | | | |
| Participant variance 0.204 |  |  |  |  |
| N participant 20, 180 Observations in total |  |  |  |  |
| Marginal $R^2$ / Conditional $R^2$ 0.019 / 0.901 | | | | |

**Note:** The three-way interaction did not contribute significantly to the model ( $\chi^2(2) = 0.565$ , $p = 0.754$ ), so it was removed; the same was the case for attentional focus by context ( $\chi^2(2) = 0.581$ , $p = 0.748$ ) and attentional focus by colour ( $\chi^2(1) = 0.396$ , $p = 0.529$ ).
This model violated the assumption of normally distributed residuals, suffering from excessive outliers. To facilitate interpretation, we report estimates and confidence intervals from a bootstrapped model (with 1000 samples), obtained using the *lmeresampler* R package. We report p values and other properties of the model (e.g., residual variance and r squared) from the model fitted to log transformed amplitudes, which is compliant with all the assumptions and exhibits the same statistically significant effects.

**Table S8.** *LMM -* SSVEP amplitudes elicited by blue. Model equation: amplitude ∼ 1 + (1|participant)

|  | Amplitude |  |  |  |
| --- | --- | --- | --- | --- |
| *Predictors* | *Estimate* | *95% CI* | *t* | *p* |
| (Intercept) | 1.075 | 0.912 –<br>1.237 | 13.59 | <.001 |
| *Random effects* |  |  |  |  |
| Residual variance, $\sigma^2 = 0.011$ | | | | |
| Participant variance 0.121 |  |  |  |  |
| ICC 0.92 |  |  |  |  |
| N participant 20, 60 Observations in total |  |  |  |  |
| Marginal $R^2$ / Conditional $R^2$ 0 / 0.919 | | | | |

**Note:** Attentional focus did not contribute significantly to the model ( $\chi^2(2) = 2.712$ , $p = 0.258$ ).

**Table S9.**
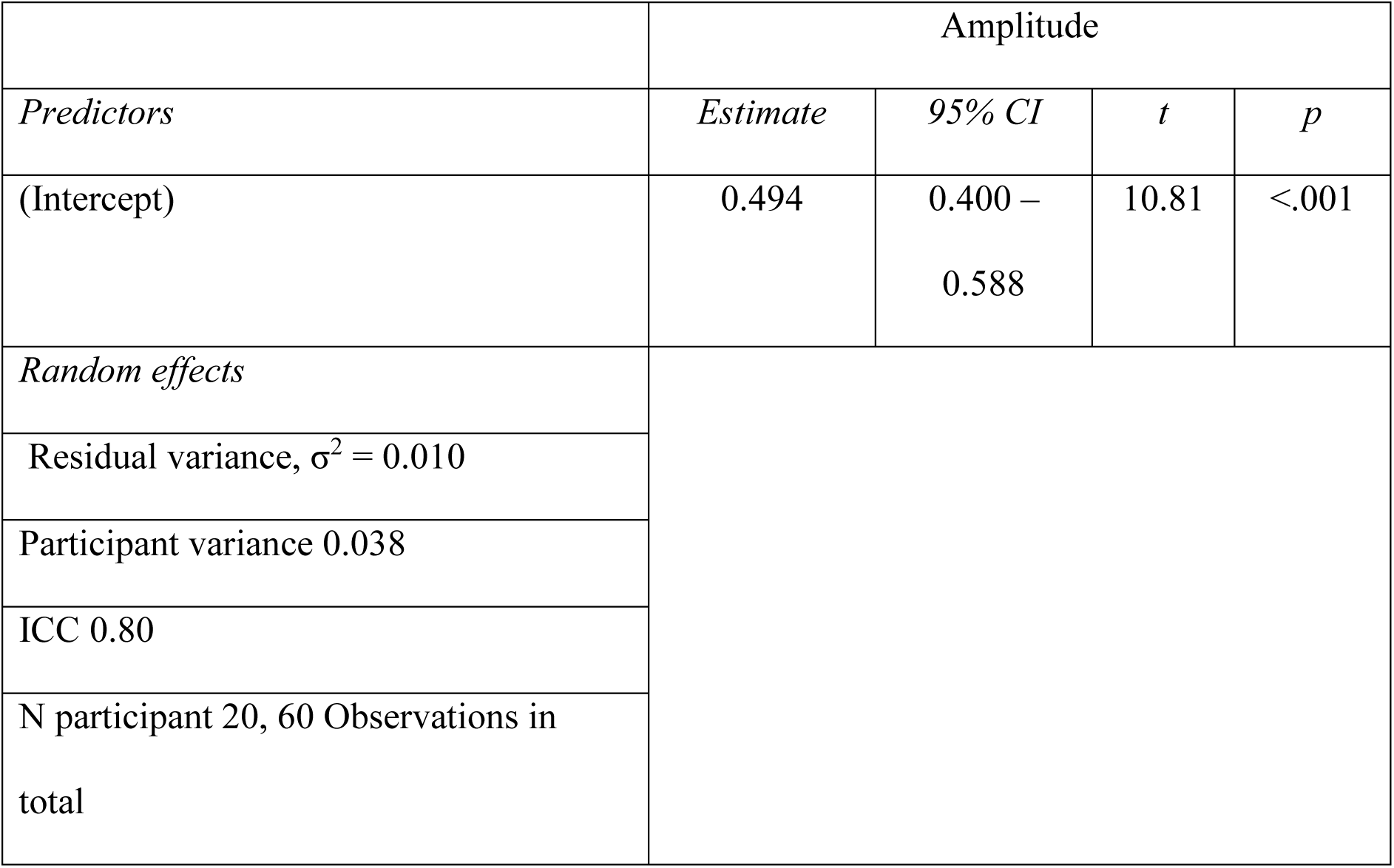
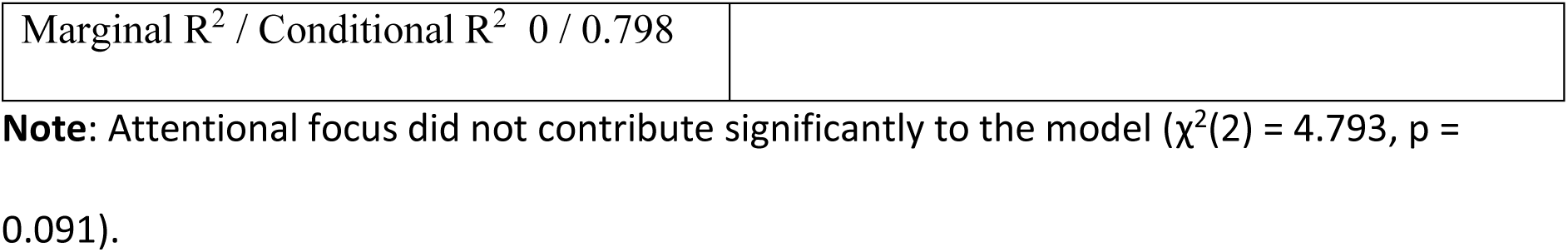
*LMM -* SSVEP amplitudes elicited by lime. Model equation: amplitude ∼ 1 + (1|participant)

|  | Amplitude |  |  |  |
| --- | --- | --- | --- | --- |
| *Predictors* | *Estimate* | *95% CI* | *t* | *p* |
| (Intercept) | 0.494 | 0.400 –<br>0.588 | 10.81 | <.001 |
| *Random effects* |  |  |  |  |
| Residual variance, $\sigma^2 = 0.010$ | | | | |
| Participant variance 0.038 |  |  |  |  |
| ICC 0.80 |  |  |  |  |
| N participant 20, 60 Observations in total |  |  |  |  |

|  |  |
| --- | --- |
| Marginal $R^2$ / Conditional $R^2$ 0 / 0.798 | |

**Note:** Attentional focus did not contribute significantly to the model ( $\chi^2(2) = 4.793$ , $p = 0.091$ ).

**Table S10.**
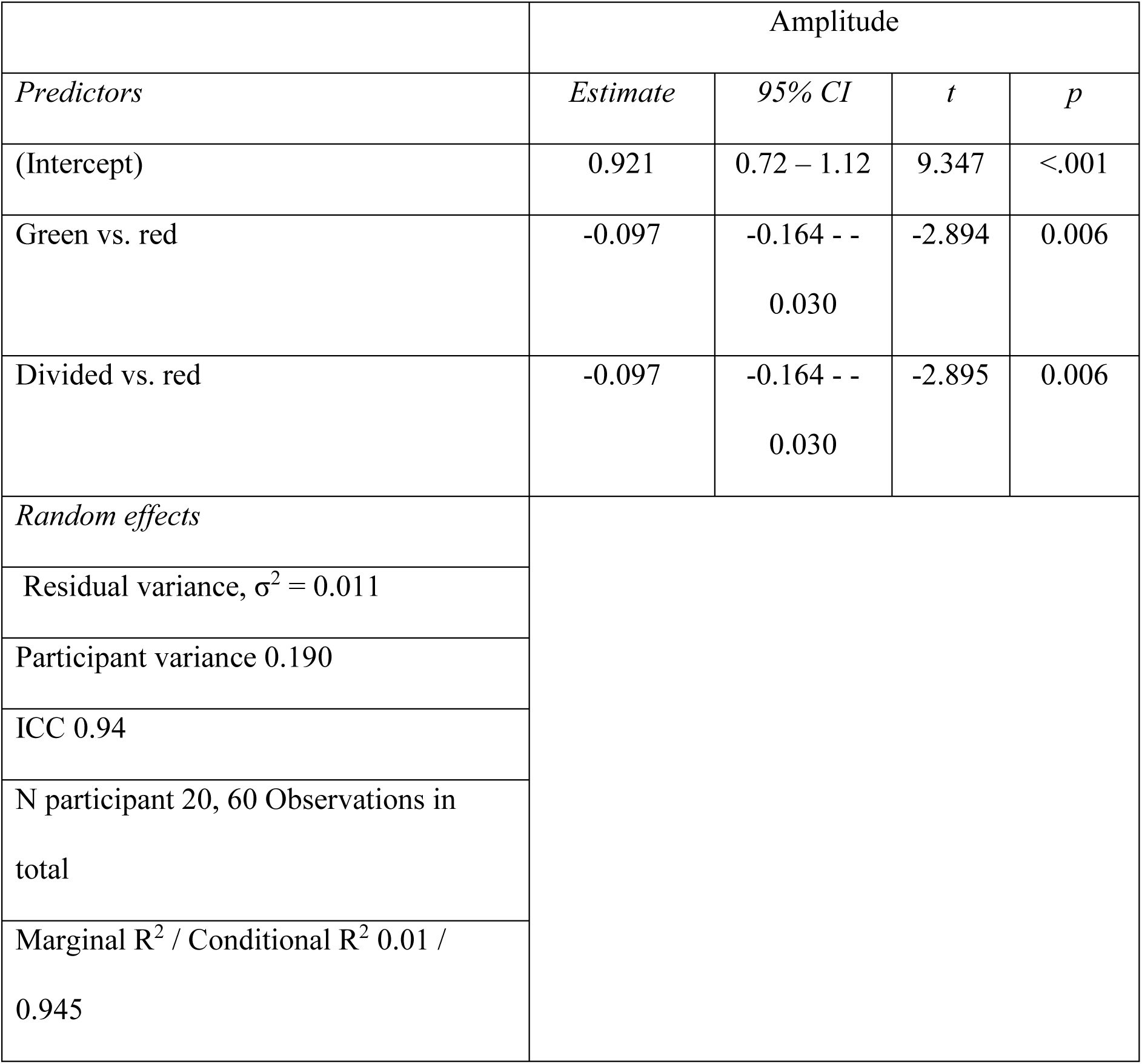
*LMM -* SSVEP amplitudes elicited by orange. Reference level was set to red focus. Model equation: amplitude ∼ 1 + attentional_focus + (1|participant)

## References

Adamian N, Andersen SK. 2022. Attentional Enhancement of Tracked Stimuli in Early Visual Cortex Has Limited Capacity. J Neurosci 42:8709–8715.

Adamian N, Andersen SK. 2023. Attentional Modulation in Early Visual Cortex: A Focused Reanalysis of Steady-state Visual Evoked Potential Studies. J Cogn Neurosci:1–25.

Adamian N, Slaustaite E, Andersen SK. 2019. Top–down attention is limited within but not between feature dimensions. J Cogn Neurosci 31:1173–1183.

Allard R, Faubert J. 2014. An expansive, cone-specific nonlinearity enabling the luminance motion system to process color-defined motion. J Vis 14:2–2.

Andersen SK, Fuchs S, Muller MM. 2011. Effects of Feature-selective and Spatial Attention at Different Stages of Visual Processing. J Cogn Neurosci 23:238–246.

Andersen SK, Hillyard SA, Muller MM. 2008. Attention facilitates multiple stimulus features in parallel in human visual cortex. Curr Biol 18:1006–1009.

Andersen SK, Hillyard SA, Muller MM. 2013. Global Facilitation of Attended Features Is Obligatory and Restricts Divided Attention. J Neurosci 33:18200–18207.

Andersen SK, Muller MM. 2010. Behavioral performance follows the time course of neural facilitation and suppression during cued shifts of feature-selective attention. Proc Natl Acad Sci U S A 107:13878–13882.

Andersen SK, Muller MM, Hillyard SA. 2015. Attentional Selection of Feature Conjunctions Is Accomplished by Parallel and Independent Selection of Single Features. J Neurosci 35:9912–9919.

Andersen SK, Müller MM, Hillyard SA. 2011. Tracking the allocation of attention in visual scenes with steady-state evoked potentials. In: Posner MIC, editor. Cognitive Neuroscience of Attention (2nd ed.) New York: Guilford.

Andersen SK, Muller MM, Martinovic J. 2012. Bottom-up biases in feature-selective attention. J Neurosci 32:16953–16958.

Bae GY, Olkkonen M, Allred SR, Flombaum JI. 2015. Why Some Colors Appear More Memorable Than Others: A Model Combining Categories and Particulars in Color Working Memory. J Exp Psychol-Gen 144:744–763.

Bays PM, Catalao RFG, Husain M. 2009. The precision of visual working memory is set by allocation of a shared resource. J Vis 9:7.1–11.

Becker SI. 2010. The role of target–distractor relationships in guiding attention and the eyes in visual search. J Exp Psychol Gen 139:247–265.

Boylan MR, Panitz C, Tebbe A-L, Vieweg P, Forschack N, Müller MM, Keil A. 2023. Feature-based Attentional Amplitude Modulations of the Steady-state Visual Evoked Potentials Reflect Blood Oxygen Level Dependent Changes in Feature-sensitive Visual Areas. J Cogn Neurosci:1–15.

Brouwer GJ, Heeger DJ. 2009. Decoding and Reconstructing Color from Responses in Human Visual Cortex. J Neurosci 29:13992–14003.

Brouwer GJ, Heeger DJ. 2013. Categorical Clustering of the Neural Representation of Color. J Neurosci 33:15454–15465.

Carrasco M. 2014. Spatial Covert Attention: Perceptual Modulation. In: Nobre AC, Kastner S, editors. The Oxford Handbook of Attention Oxford: Oxford University Press p 184–230.

Champely S, Ekstrom C, Dalgaard P, Gill J, Weibelzahl S, Anandkumar A, Ford C, Volcic R, De Rosario H. 2020. Power analysis functions along the lines of Cohen (1988), version 1.3. In, 17.3.2020. ed: https://github.com/heliosdrm/pwr.

Duncan J. 1984. Selective attention and the organization of visual information. J Exp Psychol Gen 113:501–517.

Eriksen CW, St. James JD. 1986. Visual attention within and around the field of focal attention. Perception & Psychophysics 45:225–240.

Fairchild MD. 1996. Refinement of the RLAB color space. Color Research & Application 21:338–346.

Fairchild MD. 2013. Colour Appearance Models (3rd revised edition): Wiley-Blackwell.

Forsberg A, Johnson W, Logie RH. 2019. Aging and feature-binding in visual working memory: The role of verbal rehearsal. Psychology and aging.

Found A, Muller HJ. 1996. Searching for unknown feature targets on more than one dimension: Investigating a ’’dimension-weighting’’ account. Perception & Psychophysics 58:88–101.

Geng JJ, Witkowski P. 2019. Template-to-distractor distinctiveness regulates visual search efficiency. Curr Opin Psychol 29:119–125.

Gundlach C, Forschack N, Müller MM. 2023. Global attentional selection of visual features is not associated with selective modulation of posterior alpha-band activity. Psychophysiology 60:e14244.

Gundlach C, Wehle S, Müller MM. 2023. Early sensory gain control is dominated by obligatory and global feature-based attention in top-down shifts of combined spatial and feature-based attention. Cereb Cortex 33:10286–10302.

Hartig F. 2022. DHARMa: Residual Diagnostics for Hierarchical (Multi-Level/Mixed) Regression Models. Version R package version 0.4.6.

Jaeger TF. 2008. Categorical Data Analysis: Away from ANOVAs (transformation or not) and towards Logit Mixed Models. J Mem Lang 59:434–446.

Liesefeld HR, Müller HJ. 2019. Distractor handling via dimension weighting. Current Opinion in Psychology 29:160–167.

Liu T. 2019. Feature-based attention: effects and control. Current Opinion in Psychology 29:187–192.

Liu T, Jigo M. 2017. Limits in feature-based attention to multiple colors. Atten Percept Psychophys 79:2327–2337.

Loy A. 2023. lmeresampler: Bootstrap Methods for Nested Linear Mixed-Effects Models. Version 0.2.4.

Lüdecke D, Ben-Shachar MS, Patil I, Waggoner P, Makowski D. 2021. performance: An R Package for Assessment, Comparison and Testing of Statistical Models. Journal of Open Source Software 6:3139.

Macmillan NA, Creelman CD. 2004. Detection Theory: A user’s guide: Psychology Press.

Malkoc G, Kay P, Webster MA. 2005. Variations in normal color vision. IV. Binary hues and hue scaling. Journal of the Optical Society of America A 22:2154–2168.

Martinez-Trujillo JC, Treue S. 2004. Feature-based attention increases the selectivity of population responses in primate visual cortex. Curr Biol 14:744–751.

Martinovic J, Andersen SK. 2018. Cortical summation and attentional modulation of combined chromatic and luminance signals. Neuroimage 176:390–403.

Martinovic J, Hardman A. 2016. Color and Visual Search, Color Singletons. In: Luo MR, editor. Encyclopedia of Color Science and Technology New York: Springer.

Martinovic J, Meyer G, Muller MM, Wuerger SM. 2009. S-cone signals invisible to the motion system can improve motion extraction via grouping by color. Vis Neurosci 26:237–248.

Martinovic J, Paramei GV, MacInnes WJ. 2020. Russian blues reveal the limits of language influencing colour discrimination. Cognition 201:104281.

Martinovic J, Wuerger SM, Hillyard SA, Müller MM, Andersen SK. 2018. Neural mechanisms of divided feature-selective attention to colour. NeuroImage 181:670–682.

Matera CN, Emery KJ, Volbrecht VJ, Vemuri K, Kay P, Webster MA. 2020. Comparison of two methods of hue scaling. Journal of the Optical Society of America A, Optics, image science, and vision 37:A44–A54.

Mullen KT, Yoshizawa T, Baker J, Curtis L. 2003. Luminance mechanisms mediate the motion of red-green isoluminant gratings: the role of “temporal chromatic aberration”. Vision Res 43:1237–1249.

Nakayama K, Silverman GH. 1984. Temporal and spatial characteristics of the upper displacement limit for motion in radom dots. Vision Res 24:293–299.

Ort E, Olivers CNL. 2020. The capacity of multiple-target search. Vis Cogn 28:330–355.

Oxner M, Martinovic J, Forschack N, Lempe R, Gundlach C, Müller M. 2023. Global enhancement of target color—not proactive suppression—explains attentional deployment during visual search. J Exp Psychol Gen 152:1705–1722.

Parker AJ. 2020. Intermediate level cortical areas and the multiple roles of area V4. Current Opinion in Physiology 16:61–67.

Posner MIC, Snyder RR, Davidson BJ. 1980. Attention and detection of signals. J Exp Psychol Gen 109:160–174.

R_Core_Team. 2016. R: A language and environment for statisticall computing. Vienna, Austria: R Foundation for Statistical Computing.

Regan BC, Reffin JP, Mollon JD. 1994. Luminance noise and the rapid determination of discrimination ellipses in color deficiency Vision Res 34:1279–1299.

Schielzeth H, Dingemanse NJ, Nakagawa S, Westneat DF, Allegue H, Teplitsky C, Réale D, Dochtermann NA, Garamszegi LZ, Araya-Ajoy YG. 2020. Robustness of linear mixed-effects models to violations of distributional assumptions. Methods in Ecology and Evolution 11:1141–1152.

Schiller F, Valsecchi M, Gegenfurtner KR. 2018. An evaluation of different measures of color saturation. Vision Res 151:117–134.

Scolari M, Byers A, Serences JT. 2012. Optimal Deployment of Attentional Gain during Fine Discriminations. The Journal of Neuroscience 32:7723.

Scolari M, Serences JT. 2009. Adaptive Allocation of Attentional Gain. The Journal of Neuroscience 29:11933.

Seymour J. 2020. Why does the CIELAB a* axis point toward magenta instead of red? Color Research & Application 45:1040–1054.

Stormer VS, Alvarez GA. 2014. Feature-Based Attention Elicits Surround Suppression in Feature Space. Curr Biol 24:1985–1988.

Thayer DD, Sprague TC. 2023. Feature-Specific Salience Maps in Human Cortex. J Neurosci 43:8785–8800.

Toffanin P, de Jong R, Johnson A, Martens S. 2009. Using frequency tagging to quantify attentional deployment in a visual divided attention task. Int J Psychophysiol 72:289–298.

Treue S, Martinez-Trujillo JC. 1999. Feature-based attention influences motion processing gain in macaque visual cortex. Nature 399:575–579.

Verghese P, Kim YJ, Wade AR. 2012. Attention Selects Informative Neural Populations in Human V1. J Neurosci 32:16379–16390.

Wickens TD. 2002. Elementary signal detection theory. New York, NY, US: Oxford University Press.

Wickham H, Averick M, Bryan J, Chang W, McGowan L, Francois R, Grolemund G, Hayes A, Henry L, Hester J, Kuhn M, Pedersen T, Miller E, Bache S, Mueller K, Ooms J, Robinson D, Seidel D, Spinu V, Takahashi K, Vaughan D, Wilke C, Woo K, Yutani H. 2019. Welcome to the tidyverse. Journal of Open Source Software 4:1686.

Wolfe JM, Van Wert MJ. 2010. Varying Target Prevalence Reveals Two Dissociable Decision Criteria in Visual Search. Curr Biol 20:121–124.

Wuerger SM, Ruppertsberg A, Malek S, Bertamini M, Martinovic J. 2011. The integration of local chromatic motion signals is sensitive to contrast polarity. Vis Neurosci 28:239–246.

Yu X, Zhou Z, Becker SI, Boettcher SEP, Geng JJ. 2023. Good-enough attentional guidance. Trends Cogn Sci 27:391–403.

